# Adaptive thermogenesis in mice requires adipocyte light-sensing via Opsin 3

**DOI:** 10.1101/721381

**Authors:** Gowri Nayak, Shruti Vemaraju, Kevin X. Zhang, Yoshinobu Odaka, Ethan D. Buhr, Amanda Holt-Jones, April N. Smith, Brian A. Upton, Jesse J. Zhan, Nicolás Diaz, Kazutoshi Murakami, Shane D’Souza, Minh-Thanh Nguyen, Shannon A. Gordon, Gang Wu, Robert Schmidt, Xue Mei, Nathan T. Petts, Matthew Batie, Sujata Rao, Takahisa Nakamura, Alison M. Sweeney, John B. Hogenesch, Russell N. Van Gelder, Joan Sanchez-Gurmaches, Richard A. Lang

## Abstract

Almost all life forms can decode light information for adaptive advantage. Examples include the visual system, where photoreceptor signals are interpreted as images, and the circadian system, where light entrains a physiological clock. Here we describe a local, non-visual light response in mice that employs encephalopsin (OPN3, a 480 nm, blue light responsive opsin) to regulate the function of adipocytes. Germ line null and adipocyte-specific conditional null mice show a deficit in thermogenesis that is phenocopied in mice under blue-light deficient conditions. We show that blue light stimulation of adipocytes activates hormone sensitive lipase, the rate limiting enzyme in the lipolysis pathway, and that this is OPN3-dependent. *Opn3* adipocyte conditional null mice also use reduced levels of fat mass when fasted and cold exposed further suggesting a lipolysis deficit. These data suggest the hypothesis that in mice, a local, OPN3-dependent light response in adipocytes is a mechanism for regulation of energy homeostasis.

## Introduction

The detection of photons by animals has been exploited for adaptive advantage in many different ways. The visual sense – irradiance detection by photoreceptors in the retina and the perception of images in the brain – is the most obvious example because it is a component of our conscious existence. However, most animals have a parallel system of non-visual ocular photoreceptors. In mammals, the best characterized are the melanopsin (OPN4) and neuropsin (OPN5)-expressing retinal ganglion cells that function in negative phototaxis (Johnson et al., 2010), retinal and corneal circadian clock entrainment (Buhr et al., 2015), the pupillary light reflex (Hattar et al., 2002) and in eye development (Nguyen et al., 2019; Rao et al., 2013).

Extraocular photoreceptors are found throughout the animal kingdom. They exist within the skin of frogs (Moriya et al., 1996; Provencio et al., 1998), within pineal organs that produce melatonin (Okano et al., 1994), and as deep brain photoreceptors that regulate seasonal breeding responses in avian species (Nakane et al., 2010). Until recently, it was thought that extraocular photoreception was absent from mammals. However, ongoing characterization of *Opn3, Opn4* and *Opn5* has demonstrated expression domains outside the eye. Coupled with studies showing light-dependent signaling by these opsins (Kato et al., 2016; Kojima et al., 2011; Koyanagi et al., 2013a; Yamashita et al., 2010), this has raised the possibility of extraocular photoreception in mammals. Evidence for the function of these signals is limited to date but includes a possible role for OPN4 in acutely-regulated dilation of blood vessels (Sikka et al., 2014, 2016) and in the modulation of melanocyte pigmentation (de Assis et al., 2018; Ozdeslik et al., 2019). It has also been suggested that adipocyte function might be modulated by light stimulation of OPN4 (Ondrusova et al., 2017).

Mammals employ three different types of adipocyte for distinct functions (Giralt and Villarroya, 2013). White adipose tissue (WAT) consists primarily of white adipocytes and is the major energy storage site. Under conditions of cold exposure, WAT can differentiate to generate so called “brite” (brown-in-white) adipocytes that have a limited thermogenesis capacity (Rosenwald et al., 2013; Shabalina et al., 2013; Wu et al., 2012). Brown adipose tissue (BAT) is made up exclusively of brown adipocytes; it has a limited storage function but a key role in generating heat via non-shivering thermogenesis (Giralt and Villarroya, 2013). The process of lipolysis releases free fatty acids (FFA) from WAT and these are used systemically for cellular energy (Zechner et al., 2012). BAT uses FFA for the generation of heat via thermogenesis pathways (Ikeda et al., 2017; Kazak et al., 2015, 2017), one of which employs uncoupling protein 1 (UCP1) to separate oxidative metabolism from ATP production (Betz and Enerbäck, 2018). Thus, WAT and BAT both have important functions in the regulation of energy balance. Though it was originally believed that only newborn humans had significant depots of brown fat, it is now understood to be present in many adults (Cypess et al., 2009; van Marken Lichtenbelt et al., 2009; Nedergaard et al., 2007; Virtanen et al., 2009). Gathering evidence suggests that activation of BAT might be valuable in protection against metabolic syndrome (Harms and Seale, 2013; Seale, 2013).

Here we describe an extraocular function for the G-protein coupled family member and non-canonical opsin encephalopsin (OPN3)(Blackshaw and Snyder, 1999) in the light-dependent regulation of adipocyte function. When cold challenged, mice with an adipocyte-specific deletion of *Opn3* fail to defend their body temperature normally, show an attenuated induction of thermogenesis pathway genes in BAT and use less fat mass. These phenotypes are reproduced in mice that are raised in the absence of the blue light wavelengths that normally stimulate OPN3. These metabolic perturbations appear to be explained by the OPN3- and blue-light dependence of the lipolysis response, a pathway that normally provides fatty acids as fuel for thermogenesis. These data identify an extraocular mechanism for light information decoding that regulates energy homeostasis.

## Results

### *Opsin 3* is expressed in adipocytes

An initial survey of *Opn3* expression in mice revealed that adipose tissue was positive. No reliable antibodies for murine OPN3 are presently available and so to assess expression we took advantage of three alleles (Fig. S1), *Opn3*^*lacz*^, *Opn3*^*cre*^ (in combination with the tdTomato reporter *Ai14*) and *Opn3-eGFP*, an expression reporter transgene based on a bacterial artificial chromosome (GENSAT #030727-UCD). The interscapular adipose tissue (iAT) depot comprises interscapular subcutaneous white adipose tissue (iscWAT) and interscapular brown adipose tissue (iBAT). Xgal labeling of control cryosections from P16 iAT showed no background labeling in wild type (Fig. 1A) but intense labeling in the *Opn3*^*lacz/lacz*^ iscWAT (Fig. 1B). Labeled adipocytes were not detected in control iBAT (Fig. 1C). In iBAT from *Opn3*^*lacz/lacz*^, Xgal labeled cells were not readily apparent at low magnification but at higher magnification and bright transillumination, a subset of expressing brown adipocytes was detected (Fig. 1D).

**Figure 1.**
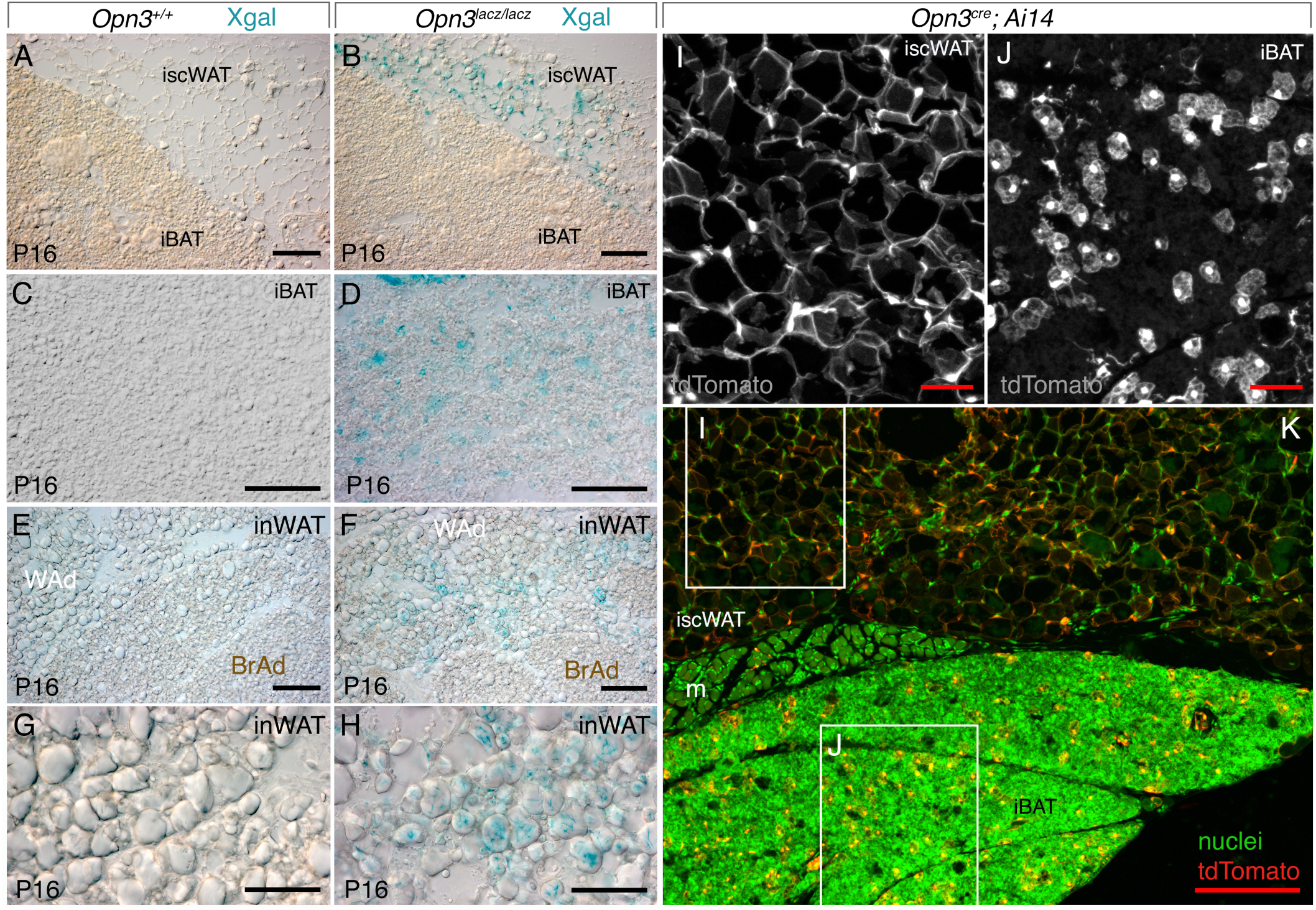
Expression of *Opn3* in iAT and inWAT. (A-D) Xgal labeled wild type (A, C) and *Opn3*^*lacz/lacz*^ (B, D) cryosections of interscapular adipose tissue (iAT) including interscapular subcutaneous white adipose tissue (A, B, iscWAT) and interscapular brown adipose tissue (A-D, iBAT) at P16. (E-H) Xgal labeled wild type (E, G) and *Opn3*^*lacz/lacz*^ (F, H) cryosections of inguinal white adipose (inWAT) tissue including white (WAd) and “brite” (BrAd) adipocytes. (I-K) Detection of tdTomato (red, greyscale) in *Opn3*^*cre*^; *Ai14* mice for iAT showing positive cells in iscWAT (I) and iBAT (J). iscWAT and iBAT are separated by a leaflet of muscle (m) that is visible in some sections. Labeling of nuclei with Hoechst33258 is presented in green. In (A,B,E,F,K) scale bars are 100 µm. In (C,D,G,H,I,J) scale bars are 50 µm.

Neonatal inWAT has a high content of “brite” adipocytes (Fig. 1E, F, BrAd). In P16 inguinal WAT (inWAT) from control mice, neither the large unilocular white or the smaller brite adipocytes were Xgal-labeled (Fig. 1E, G). By contrast, the large, unilocular white adipocytes from *Opn3*^*lacz/lacz*^ mice were Xgal positive (Fig. 1F, H). When we used the *Opn3*^*cre*^ allele (Fig. S1) to convert the tdTomato reporter *Ai14* (Fig. 1I-K) cryosections showed that almost all adipocytes within the iscWAT were positive (Fig. 1I, K). In iBAT, a subset of brown adipocytes was positive (Fig. 1J, K). In an assessment of inWAT, iscWAT and iBAT, the *Opn3-eGFP* reporter confirmed expression of *Opn3* in the majority of white adipocytes and a subset of brown adipocytes in iBAT (Fig. S2). Finally, the GTEx database showed that in human subcutaneous and omental adipose tissue, OPN3 was expressed at low-to-moderate levels of 2.9 and 3.1 tags per million, respectively (Fig. S2). These data indicate that both mouse and human adipose tissues express *Opn3*. Expression of *Opn3* in adipocytes raised the possibility that adipose tissue might be directly light responsive.

### Photon flux within iscWAT and iBAT is sufficient for opsin activation

If OPN3 in adipocytes is to mediate light responsiveness, there must sufficient photon flux within adipose tissue to activate this opsin. To measure this, we designed an experiment in which the photon flux from a light source of known irradiance could be measured directly within adipose tissue using an optic fiber-based microprobe (Fig. 2A). The Holt-Sweeney microprobe (HSM, Fig. 2B) is fabricated from a light-shielded optic fiber but has a scattering spherical diffusing tip so that it can accept photons with equal probability from all directions (it is a single cell-sized detector of scalar irradiance)(Holt et al., 2014). The microprobe is mounted within a pulled Pasteur pipette for structural stability and a stereotaxic frame allows depth of tissue penetration to be precisely controlled. The microprobe tip is small enough (∼60 μm) to allow adipose tissue penetration after a skin incision has been made. Measurements of photon flux at the surface of iAT and at depths of 0.5-2.5 mm (Fig. 2C) were made over the 350-800 nm spectral range. At the λ_max_ of 480 nm for OPN3 (Fig. 2D), the measured photon flux was 5×10^14^ photons cm^−2^sec^−1^ at 0.5 mm (approximately the deepest point within the iscWAT, Fig. 2C) and 2×10^13^ photons cm^−2^sec^−1^ at 2.5 mm (the deepest point within the iBAT, Fig. 2C). These intra-tissue photon fluxes were obtained with a surface illumination (2×10^15^ photons cm^−2^sec^−1^) that represents approximately 1% of clear sky sunlight intensity. Therefore, the light attenuation ranged from less than one log quanta at 0.5 mm to just over two log quanta at 2.5 mm relative to irradiance at the surface of the skin. Atypical opsins have threshold signaling responses at photon fluxes as low as 10^10^ photons cm^−2^sec^−1^ (Wong, 2012). These data thus indicate that the photon flux within iscWAT and iBAT is easily sufficient for opsin activation and signaling.

**Figure 2.**
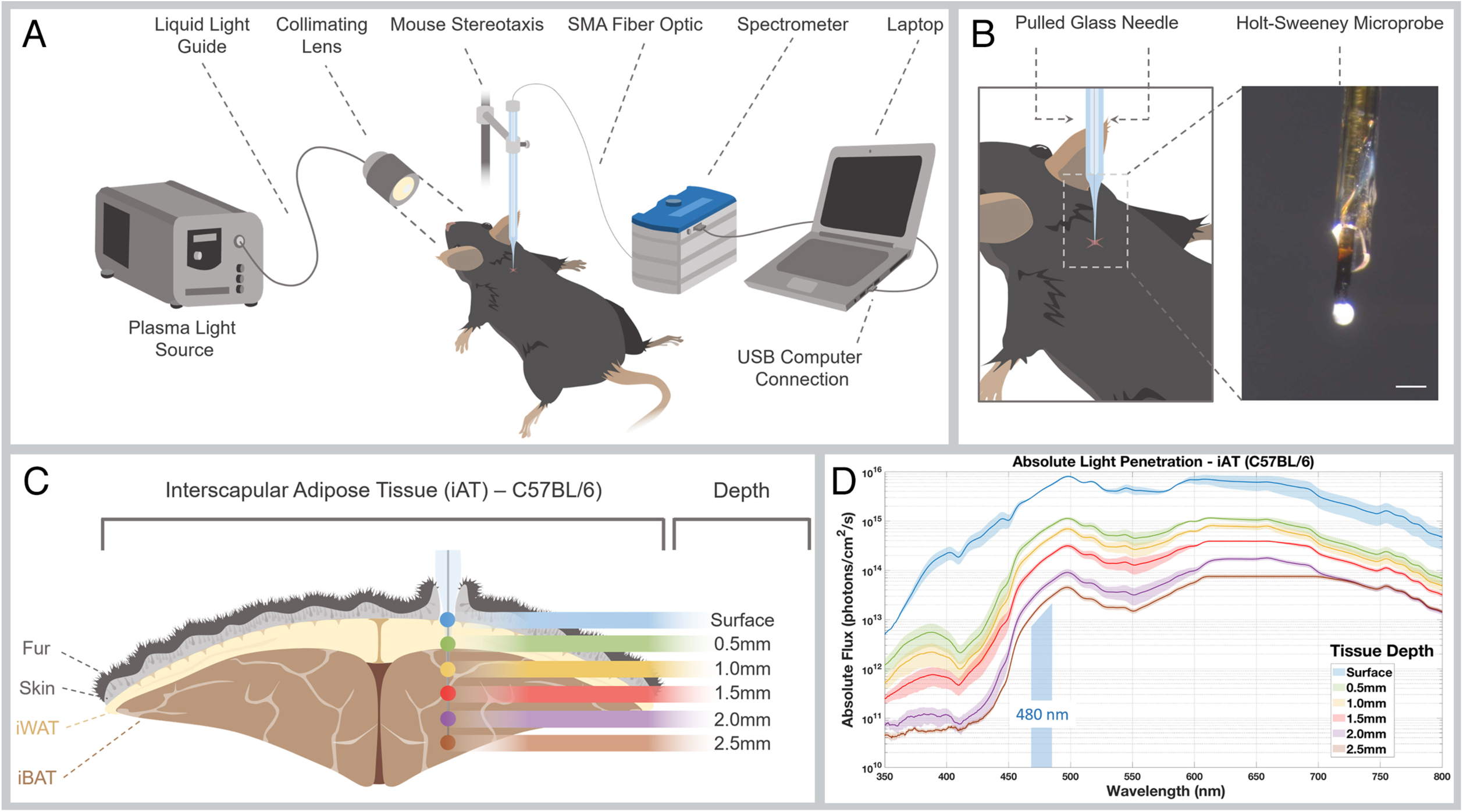
Measurement of photon flux within interscapular brown and subcutaneous white adipose tissue. (A) Schematic describing experimental set up for measuring intra-tissue photon flux. Photons are produced by a plasma light source and *via* a collimating lens are directed towards an anesthetized mouse into which the optic fiber measurement probe is placed using a stereotaxic frame. Photons are detected by an OceanOptics spectrometer controlled by a laptop computer. (B) The Holt-Sweeney microprobe (scale bar is 100 μm) probe is an optic fiber with a transparent spherical tip that accepts photons over approximately 4π steradians. (C) Measurements were taken with the microprobe at various depths in the interscapular adipose tissue. 0.5 mm corresponds to subcutaneous white adipose tissue. 1.0-2.5 mm is within the interscapular brown adipose tissue. (D) Absolute photon flux from 350-800 nm color-coded for depth of probe penetration within interscapular adipose tissue. The uppermost blue trace is surface flux and, at the *λ*max for OPN3, is about 2×10^15^ photons cm^−2^sec^−1^, approximately 1% of maximum sunlight intensity. At the maximum 2.5 mm depth (brown trace) the flux at the OPN3 *λ*max is approximately 10^13^ photons cm^−2^sec^−1^. Each trace is averaged data from n=3 mice, and color shading is ±SEM.

### Transcriptome analysis suggests that *Opsin 3* regulates metabolism

To address the function of OPN3, we performed a microarray-based transcriptome analysis on P16 control and *Opn3* germ-line null mice. Tissues for this analysis were harvested from neonatal mice at P16 from two tissues that express *Opn3* (iAT and inWAT) and one tissue, liver, that has very low levels. Using the AltAnalyze suite (Emig et al., 2010; Salomonis, 2012), we identified differentially regulated transcripts that fell into functional clusters and pathways. Clustering based on significant Z-scores for WikiPathway models is noted (Fig. S3A-C) but this information is also developed into a more detailed overall schematic (Fig. S3D-H). In addition, a subset of this schematic for WAT is shown in Figure 3A. In these schematics, each box represents a transcript where red and blue color coding indicates up- or down-regulation, respectively. With a few noted exceptions, all *Opn3*-dependent transcript regulation is significant to p<0.05. The *Opn3* transcript showed the highest negative fold change in *Opn3* null iAT (3.0 fold down, p=1.5×10^−4^) and inWAT (5.6 fold down, p=7.6×10^−3^) but was not significantly changed in liver where expression levels are normally very low. Overall, transcriptome analysis indicated that OPN3 activity is required for normal regulation of metabolism, as evidence by deregulation of adipocyte extracellular matrix, lipid, glucose and energy homeostasis pathways.

**Figure 3.**
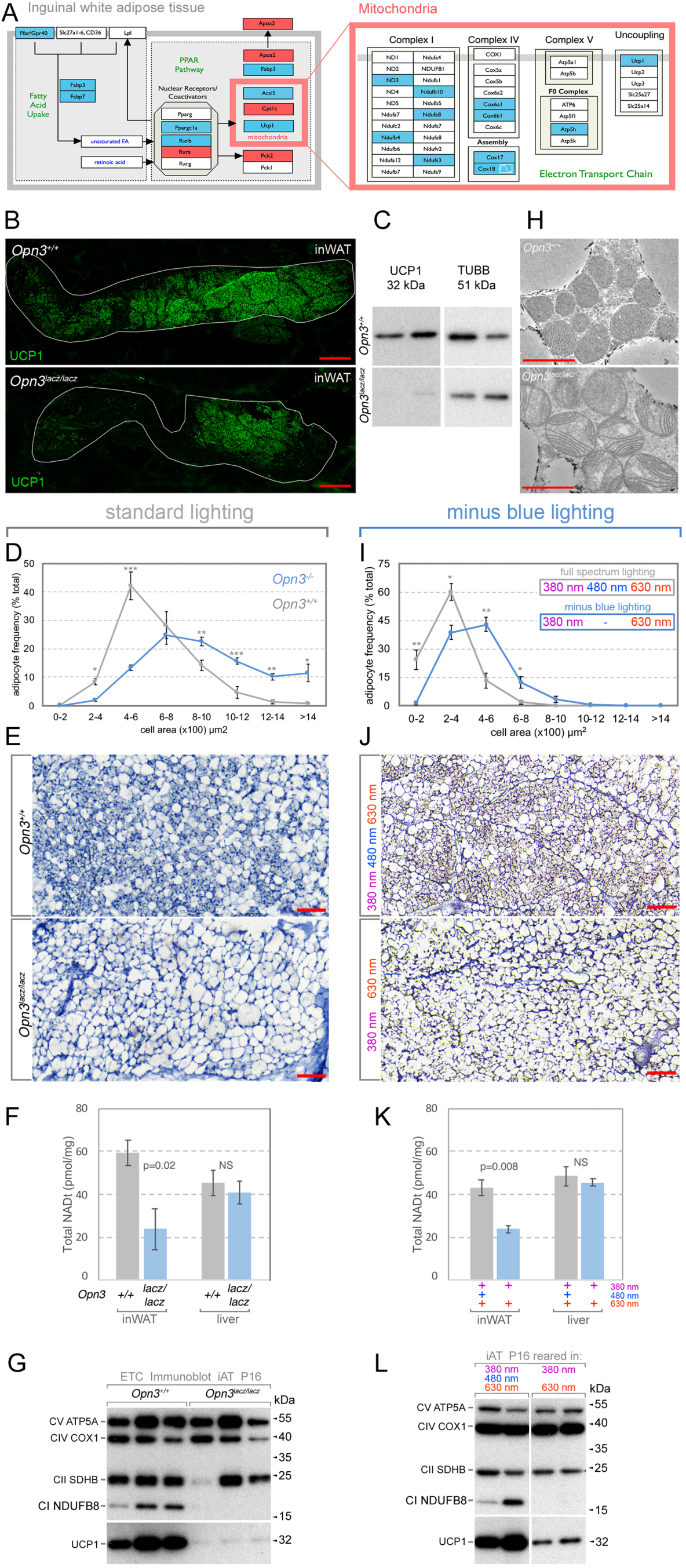
*Opn3* null and “minus blue” reared mouse adipose phenotype. (A) Schematic of clustered *Opn3*-dependent transcript changes in the PPAR and lipid uptake pathways and in the electron transport chain of mitochondria. Red and blue color-coding indicates up- and down-regulated transcripts, respectively. Schematics are modified versions of Wikipathway models for PPAR Signaling Pathway (WP2316), and for the electron Transport Chain (WP295). (B) UCP1 (green) labeling of inWAT in control and *Opn3*^*lacz/lacz*^ germ-line null animals at P16. Boundary of inWAT is outlined in white. (C) Immunoblot detecting UCP1 and β-tubulin (TUBB) in P16 inguinal WAT from *Opn3*^*+/+*^ and *Opn3*^*lacz/lacz*^ mice. UCP1 levels are low in the *Opn3* null tissue. (D, I) Adipocyte size distribution in inWAT comparing control and *Opn3*^*lacz/lacz*^ (D) and full spectrum versus “minus blue” raised mice at P16. Data are presented as mean ± s.e.m. n=3 for each genotype. Direct comparisons between genotypes at each interval were performed with Student’s T-test \**P*<0.05, \*\**P*<0.01, \*\*\**P*<0.001. (E, J) Hematoxylin staining of histological sections of P16 inWAT from *Opn3*^*+/+*^, *Opn3*^*lacz/lacz*^ (E) and full spectrum (380 nm, 480 nm, 630 nm) reared versus “minus blue” (380 nm, 630 nm) reared (J) mice. (F, K) Total NAD levels in inguinal WAT and liver for P16 *Opn3*^*+/+*^ and *Opn3*^*lacz/lacz*^ mice (F, n=4) or for mice reared either in “full spectrum” or “minus blue” lighting (K, n=3). p values calculated using Student’s T-test. In both the *Opn3* null and “minus blue” conditions, NAD level are low in inWAT. (G, L) Immunoblots detecting multiple components of the electron transport chain (ATP5A, COX1, SDHB, NDUFB8, UCP1) in P16 iAT for *Opn3*^*+/+*^ and *Opn3*^*lacz/lacz*^ (G) and “minus blue” reared mice (L). NDUFB8 and UCP1 are consistently at lower levels in *Opn3* null and “minus blue” reared mice. (H) TEM showing abnormal mitochondrial morphology in the *Opn3* null iBAT at P28. Red scale bar is 2 µm. Scale bars in red (B) 500 µm (E, J) 100 µm (H) 2 µm.

In inWAT, *Opn3*-affected transcripts cluster within the PPAR pathway and within the electron transport chain of mitochondria (Fig. 3A). The PPAR pathway regulates adipocyte size as well as lipid metabolism and energy generation (Fan and Evans, 2015; Liu et al., 2007). This pathway regulates energy generation in part because multiple components of the lipolysis pathway, including HSL (hormone sensitive lipase), ATGL (adipose triglyceride lipase)(Leone et al., 1999; Rhee et al., 2003; Vega et al., 2000) and perilipin (Arimura et al., 2004), are directly or indirectly dependent on the transcriptional co-activator PGC1a for their expression. Furthermore, the transcript for UCP1 (uncoupling protein 1) that is required for one pathway in adaptive thermogenesis (Shabalina et al., 2013) is down-regulated, presumably as a response to deregulation of its transcription factors PGC1a and RXRa/b (Puigserver et al., 1998). Notably, the transcript for lipoprotein lipase (*Lpl*), an enzyme with a role in the generation and uptake of FFA via extracellular lipolysis (Olivecrona, 2016) is deregulated in the liver of *Opn3* null mice (Fig. S3). Finally, inWAT from the *Opn3* null showed a striking cluster of 17 electron transport chain (ETC) transcripts, all of which were down-regulated (Fig. 3A). Combined, these data suggested that *Opn3* null mice might show deregulated energy metabolism.

Detection of UCP1 in inWAT either by immunofluorescence (Fig. 3B) or by immunoblot (Fig. 3C) confirmed that levels were comparatively low in the *Opn3* null. An assessment of cell size in inWAT (Fig. 3D) showed that the *Opn3* null had, on average, larger adipocytes. Hematoxylin staining of inWAT sections showed that the *Opn3* germ line null (Fig. 3E) has a lower proportion of the smaller, “brite” adipocytes, consistent with adipocyte size assessment. *Opn3* null mice do not show a change in the mass of the iAT or inWAT compared with wild type mice (Fig. S4A). We then assessed inWAT from *Opn3* null mice at P16 for total NAD content as this essential coenzyme mediator of electron transport provides a measure of mitochondrial content and function (Peek et al., 2013). Quantification showed that there was no change in liver NAD in the *Opn3* null (Fig. 3F) but that inWAT showed a reduction, consistent with reduced mitochondrial function. Though transcriptome analysis for *Opn3* null iBAT did not cluster the ETC, we were prompted by the effects of *Opn3* deficiency on inWAT to assess mitochondrial status in iBAT. Immunoblotting for the ETC components ATP5A (Complex V), COX1 (Complex IV), SDHB (Complex II), NDUFB8 (Complex I) and UCP1 revealed some variability in the presence of SDHB in the *Opn3* null but a consistently low level of both NDUFB8 and UCP1 (Fig. 3G). Consistent with these findings, at P28, mitochondrial morphology in *Opn3* null iBAT was often abnormal with a disorganized pattern of cristae (Fig. 3H).

If OPN3 is functioning in phototransductive signaling, the phenotype resulting from genetic loss of function should be mimicked by raising mice in the absence of the 480 nm blue photons that are the OPN3 ligand (Koyanagi et al., 2013b). To assess this possibility, we raised C57BL/6J mice in “minus blue” lighting conditions from just before birth and then assessed the same phenotypic characteristics that were changed in the *Opn3* null. This analysis showed that minus blue lighting resulted in inWAT with quantifiably larger adipocytes (Fig. 3I), lower brite adipocyte content by hematoxylin staining (Fig. 3J), and lower total NAD levels (Fig. 3K). Furthermore, by immunoblot, iBAT from minus blue raised mice showed low levels of NDUFB8 and lower than normal levels of UCP1 (Fig. 3L). These changes quite closely match the changes observed in the *Opn3* null raised in normal lighting (compare Fig. 3D-I, E-J, F-K and G-L) and thus support the hypothesis that OPN3 functions as a light sensor that regulates the development of adipose tissues.

### Adaptive thermogenesis in mice is promoted by blue light in an OPN3-dependent manner

In mice, body temperature is partly maintained by heat that is generated within skeletal muscle (via locomotor activity and shivering) or within BAT via non-shivering thermogenesis (NST) pathways that employ UCP1, creatine metabolism and calcium cycling (Ikeda et al., 2017; Kazak et al., 2015, 2017). The energy for thermogenesis is provided partly by the oxidative metabolism of FFA stored in adipocytes. The process of lipolysis that generates FFA is thus crucial for normal thermogenesis (Himms-Hagen, 1972). Furthermore, it has been shown that lipolysis in white adipocytes is required to fuel NST (Schreiber et al., 2017; Shin et al., 2017). Several features of the *Opn3* germ-line and “minus blue” mice suggested there might be a defect in thermogenesis: In inWAT, *Ucp1* transcripts and protein are at low levels, there is low “brite” adipocyte and NAD content and similarly, in iBAT, UCP1 and some ETC complex proteins are at low levels.

When *Opn3*^*+/+*^ and *Opn3*^*lacz/lacz*^ neonatal mice were exposed to 4°C over the course of three hours under full spectrum lighting, the core body temperature of the *Opn3* null mice was lower than normal (Fig. 4A). To take into account the possibility that this thermogenesis deficiency might be influenced by the mixed genetic background of the *Opn3* null mice (the *Opn3*^*lacz*^ allele), we generated by CRISPR a new *Opn3* loss-of-function allele on a pure C57Bl/6J background (the *Opn3*^*Δex2*^ allele) and repeated the measurement of core body temperature over a three-hour cold exposure. This showed that the effect of *Opn3* loss of function in C57Bl/6J mice was the same and resulted in a lower body temperature (Fig. 4B).

**Figure 4.**
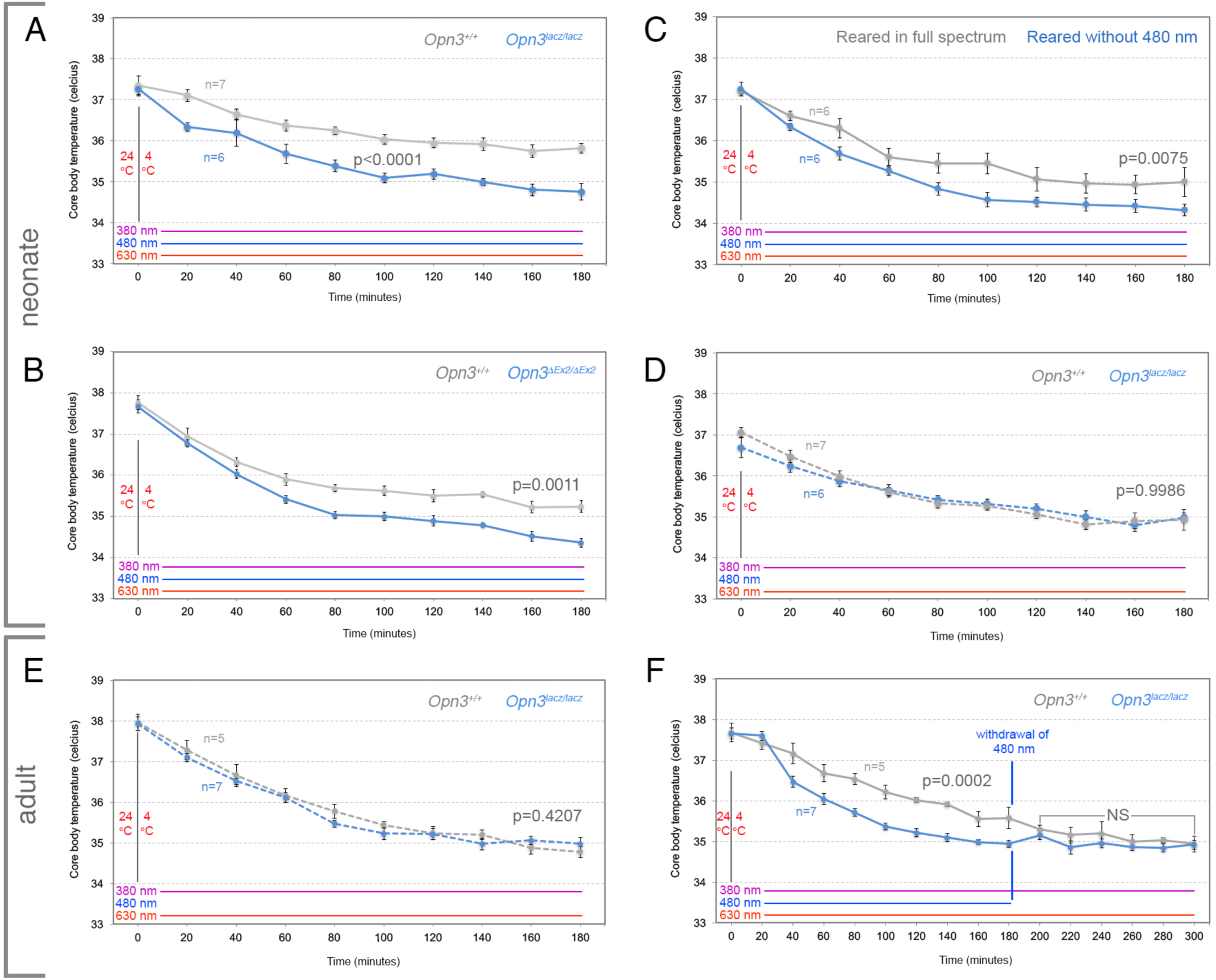
*Opn3* is required for light-dependent enhancement of the thermogenesis response. (A-F) Core body temperature (CBT) assessments over a time course after a 4°C cold exposure for P21-P24 neonatal or of adult mice of the indicated genotypes. Lighting conditions during cold exposure are indicated by the colored lines above the chart horizontal axis. (A) CBT during cold exposure in *Opn3*^*lacz/lacz*^ and control *Opn3*^*+/+*^ in full spectrum (380 nm + 480 nm + 630 nm) lighting. *Opn3*^*laczlacz*^ mice show a reduced ability to defend their body temperature. (B) CBT during cold exposure in *Opn3Δ*^*Ex2/*^*Δ*^*Ex2*^ and control *Opn3*^*+/+*^ C57BL/6J background mice in full spectrum lighting. CBT of *Opn3Δ*^*Ex2/*^*Δ*^*Ex2*^ mice is reduced compared with control. (C) CBT during cold exposure for C57BL/6J mice raised either in full spectrum (gray trace) or in minus blue (380 nm + 630 nm, blue trace) lighting. Minus blue raised mice show a reduced ability to defend their CBT. (D) CBT during cold exposure in the same cohorts of mice shown in (A) except in “minus blue” lighting. In “minus blue” *Opn3*^*+/+*^ mice show the same reduced CBT as *Opn3*^*laczlacz*^ mice in full spectrum lighting. (E) As in (D) except for adult mice. (F) CBT during cold exposure in *Opn3*^*lacz/lacz*^ and control *Opn3*^*+/+*^ in full spectrum (380 nm + 480 nm + 630 nm) lighting for 180 minutes and “minus blue” (480 nm withdrawn) lighting for a further 120 minutes. *Opn3*^*laczlacz*^ mice show reduced CBT. After withdrawal of blue light at 180 minutes, control genotype CBT descends and is then indistinguishable from the CBT of the *Opn3* null.

We could again test the assertion that OPN3 was functioning as a light sensor by assessing the thermogenesis response in mice raised in “minus blue” lighting. When C57Bl/6J mice were raised from birth in the absence of 480 nm light and assessed for core body temperature during cold exposure, they showed a lower than normal body temperature (Fig. 4C) similar to that observed in homozygotes for *Opn3*^*lacz*^ (Fig. 4A) and *Opn3*^*Δex2*^ (Fig. 4B). This showed that absence during development of the blue light that normally stimulates OPN3 could mimic genetic loss-of-function.

At the same time, it was possible that OPN3 could mediate acute light responses. We tested this possibility by performing a cold exposure assay under “minus blue” lighting but using the same cohort of *Opn3* wild type and null mice previously used under full spectrum lighting (Fig. 4A). Interestingly, under acute “minus blue” conditions, the core body temperature of control and *Opn3* null mice is statistically indistinguishable and the absolute values essentially coincident (Fig. 4D). This suggests that blue light can, via OPN3, acutely promote adaptive thermogenesis and that the activity of OPN3 fully accounts for this effect.

Neonatal mice have a beige adipocyte content that is higher than in adult mice (Sanchez-Gurmaches et al., 2012; Xue et al., 2007) and this may reflect special thermogenesis requirements given their low mass-to-surface area ratio. Thus, we were also curious as to whether adult mice showed a blue-light promoted OPN3-dependent thermogenesis response. For this analysis, we performed two types of experiment. In the first, we assessed body temperature in cohorts of adult control and *Opn3* null mice in the “minus blue” condition and showed that the body temperatures were indistinguishable (Fig. 4E). Then, with the same cohorts of mice, we repeated the assessment in full spectrum lighting and showed that over three hours of cold exposure, the core body temperature of wild type mice was higher than that on the *Opn3* null (Fig. 4F, to minute 180). This shows that, like neonatal mice, thermogenesis in adult mice is promoted by light and by OPN3 activity. After the first 180 minutes of the cold exposure experiment, we removed 480 nm stimulation and continued to assess core body temperature. We observed that over the next 20 minutes, the body temperature of wild type mice descended to a level that was indistinguishable from that of the *Opn3* null and remained at that reduced temperature for more than an hour. This provides further evidence that blue light can acutely regulate adaptive thermogenesis in an OPN3-dependent manner and that this can occur in both neonatal and adult mice.

### White adipocyte *Opn3* is required for a normal thermogenesis response

In the mouse, *Opn3* is expressed in a variety of tissues (Blackshaw and Snyder, 1999; Nissila et al., 2012; Regard et al., 2008). To determine whether we could attribute aspects of the germ line null phenotype to adipocyte *Opn3*, we performed conditional deletion of an *Opn3*^*fl*^ allele with the pan-adipocyte *Adipoq-cre* (Eguchi et al., 2011a). The iAT and inWAT of *Adipoq-cre; Opn3*^*fl/fl*^ mice did not show a significant change in mass compared with control mice (Fig. S4B) but the inWAT showed low beige content (Fig. 5A, B) and a larger adipocyte size distribution (Fig. 5C). These changes are similar to those observed in the *Opn3*^*lacz*^ homozygote null (Fig. 3D, E) and “minus blue” reared mice (Fig. 3I, J) and thus provide support for the hypothesis that OPN3 functions as a light sensor within adipocytes to regulate adipose tissue development.

**Figure 5.**
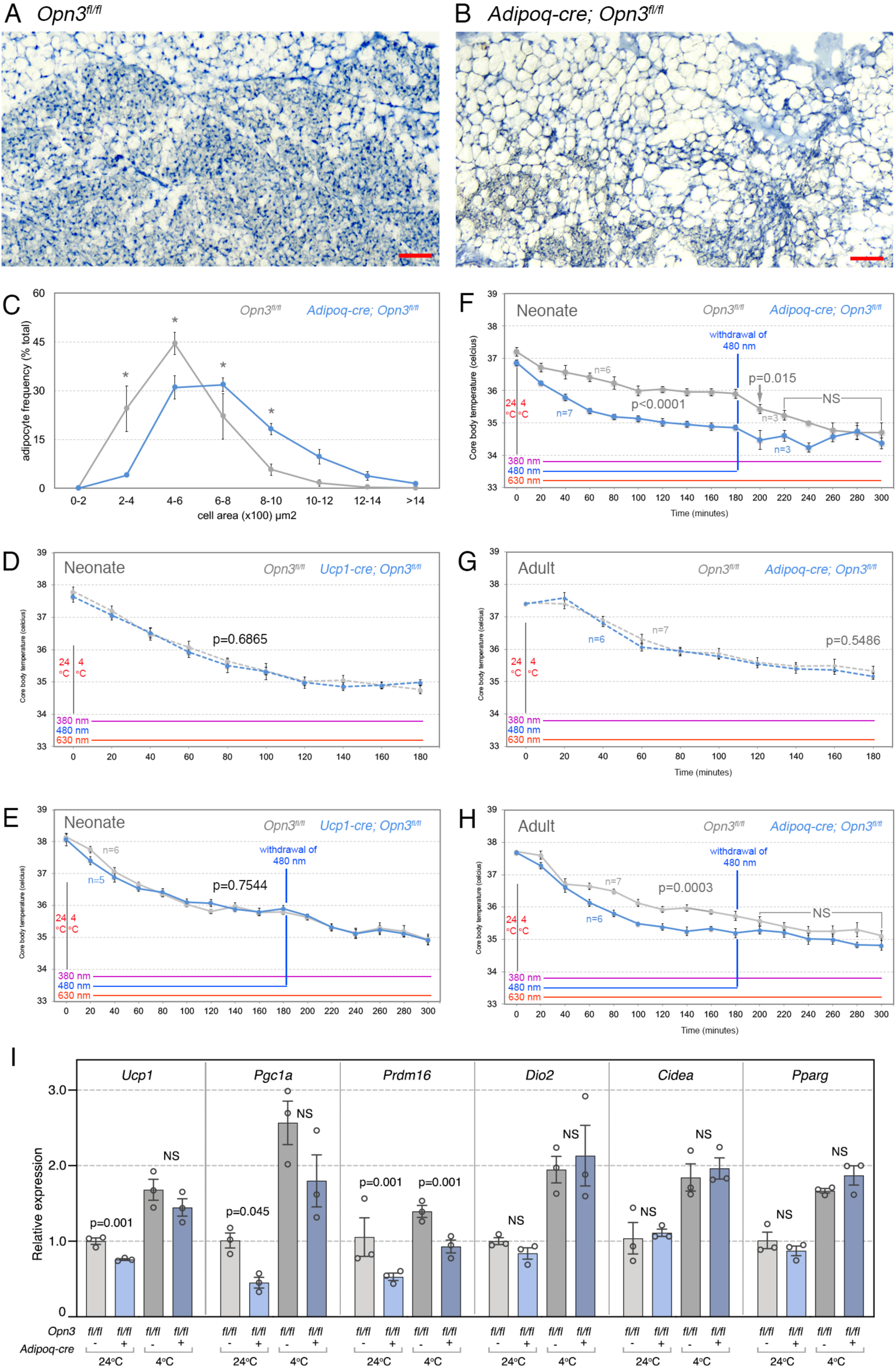
White adipocyte *Opn3* is required for a normal thermogenesis response. (A, B) Hematoxylin staining of *Opn3*^*fl/fl*^ and *Opn3*^*fl/fl*^; *Adipoq-cre* inWAT at P16. Reduced beige content of *Opn3*^*fl/fl*^; *Adipoq-cre* inWAT is apparent. (C) Adipocyte size assessment in *Opn3*^*fl/fl*^ and *Opn3*^*fl/fl*^; *Adipoq-cre* inWAT at P16. Asterisk indicates p<0.05. (D-H) Core body temperature (CBT) assessments over a time course after a 4^°C^ cold exposure for adult mice of the indicated genotypes. Lighting conditions during cold exposure are indicated by the colored lines above the chart horizontal axis. (D, E) CBT of *Opn3*^*fl/fl*^ and *Ucp1-cre; Opn3*^*fl/fl*^ mice under (D) “minus blue” conditions and (E) in full spectrum lighting for 180 minutes and then for a further 120 minutes in “minus blue”. (F) As in (E) except for cohorts of neonatal *Adipoq-cre; Opn3*^*fl/fl*^ and control *Opn3*^*fl/fl*^ mice. (G) As in (D) except for cohorts of adult *Adipoq-cre; Opn3*^*fl/fl*^ and control *Opn3*^*fl/fl*^ mice. (H) As in (F) except for adult mice. (I) Relative expression of transcripts for the thermogenesis pathway genes *Ucp1, Pgc1a, Prdm16, Dio2, Cidea* and *Pparg* in iBAT from mice of the indicated genotypes. iBAT was harvested from control mice in ambient temperature (24°C) and thos exposed to 4°C for 3 hours.

The action of both brown and white adipocytes is required for adaptive thermogenesis; brown adipocytes generate heat while white adipocytes are a major source of the FFA that are used as heating fuel (Giralt and Villarroya, 2013). Since both brown and white adipocytes express *Opn3*, it was possible that the thermogenesis deficit of the *Opn3* null mice could be explained either by a brown or white adipocyte defect (or possibly both). Thus, we measured core body temperature in cohorts of neonatal mice in which *Opn3* was conditionally deleted either in only brown adipocytes with *Ucp1-cre* (Kong et al., 2014) or in all adipocytes with *Adipoq-cre* (Eguchi et al., 2011a). For *Ucp1-cre* cohorts of mice (Fig. 5D, E), we performed cold exposure assays and measured core body temperature first over three hours in “minus blue” lighting (Fig. 5D) and then repeated the assay in full spectrum lighting with blue light withdrawal at minute 180 (Fig. 5E). In neither assay was the body temperature of *Ucp1-cre; Opn3*^*flf/fl*^ mice distinguishable from control *Opn3*^*fl/fl*^ mice. This indicated that brown adipocyte *Opn3* was not required for a normal thermogenesis response.

By contrast, neonatal *Adipoq-cre; Opn3*^*fl/fl*^ mice in full spectrum lighting showed a more limited ability than *Opn3*^*fl/fl*^ control mice to defend their body temperature (Fig. 5F). When these mice were switched to “minus blue” lighting at 180 minutes of the cold exposure experiment, the body temperature of control *Opn3*^*fl/fl*^ mice rapidly changed to the lower body temperature of the conditional null (Fig. 5F). Repetition of cold-exposure experiments in adult *Adipoq-cre; Opn3*^*fl/fl*^ null mice showed a very similar thermogenesis deficit (Fig. 5G, H) in which the absence of blue light, either throughout the cold exposure (Fig. 5G), or acutely at minute 180 (Fig. 5H) could mimic the effect of germ line *Opn3* mutation. A qPCR assessment of a set of thermogenesis pathway transcripts (*Ucp1, Pgc1a, Prdm16, Dio2, Cidea and Pparg*) showed, consistent with transcriptome analysis and the thermogenesis response deficit, that *Opn3* loss-of-function in adipocytes resulted in diminished expression of *Ucp1, Pgc1a* and *Prdm16* (Fig. 5I). Combined, these data suggest that *Opsin 3* expression in white adipocytes is required for the light-dependent component of the thermogenesis response.

### White adipocyte OPN3 is required for normal lipolysis during cold exposure

The long-standing belief that BAT lipolysis is essential for non-shivering thermogenesis has been challenged by recent analysis (Schreiber et al., 2017; Shin et al., 2017). These investigators showed that inhibiting lipolysis in BAT does not impair body temperature maintenance as long as thermogenic fuel was available through WAT lipolysis or dietary sources. We observed a similar pattern in the functional requirement for *Opn3*: Conditional deletion with *Ucp1-cre* (brown adipocytes) did not produce an abnormal thermogenesis response while deletion with *Adipoq-cre* (all adipocytes) produced mice unable to defend their body temperature normally. We therefore asked whether *Opn3* in white adipocytes was necessary to provide thermogenic fuel during exposure to cold. We used two complementary approaches. First, we performed an *in vivo* fasting experiment aimed at augmenting the use of fat reserves *via* lipolysis during cold exposure (Shin et al. 2017; Schreiber et al. 2017) and second, we determined whether white adipocytes cultured *ex vivo* showed a light- and OPN3-dependent regulation of lipolysis.

In the first experiment, cohorts of control (*Opn3*^*fl/fl*^) and experimental (*Adipoq-cre; Opn3*^*fl/fl*^) mice were either fed *ad libitum* or fasted overnight and then exposed to 4°C for 3 hours (Fig. 6A). We assessed core body temperature during the 3-hour cold exposure. This showed, as expected according to prior assessment (Fig. 5F, H), that adipocyte *Opn3* was required to maintain body temperature in fed mice (Fig. 6B, light gray and light blue traces). In fasted mice, the body temperature of both genotypes was lower still (Fig. 6B, dark gray and dark blue traces). This reflects the absence of food as an energy source. The absence of food as an energy source and the absence of adipocyte *Opn3* produced a dramatic exacerbation of the thermogenesis deficit (the fasted, *Adipoq-cre; Opn3*^*fl/fl*^ mice, Fig. 6B, dark blue trace). This was consistent with the suggestion that adipocyte *Opn3* might be required for the mobilization of lipid energy stores.

**Figure 6.**
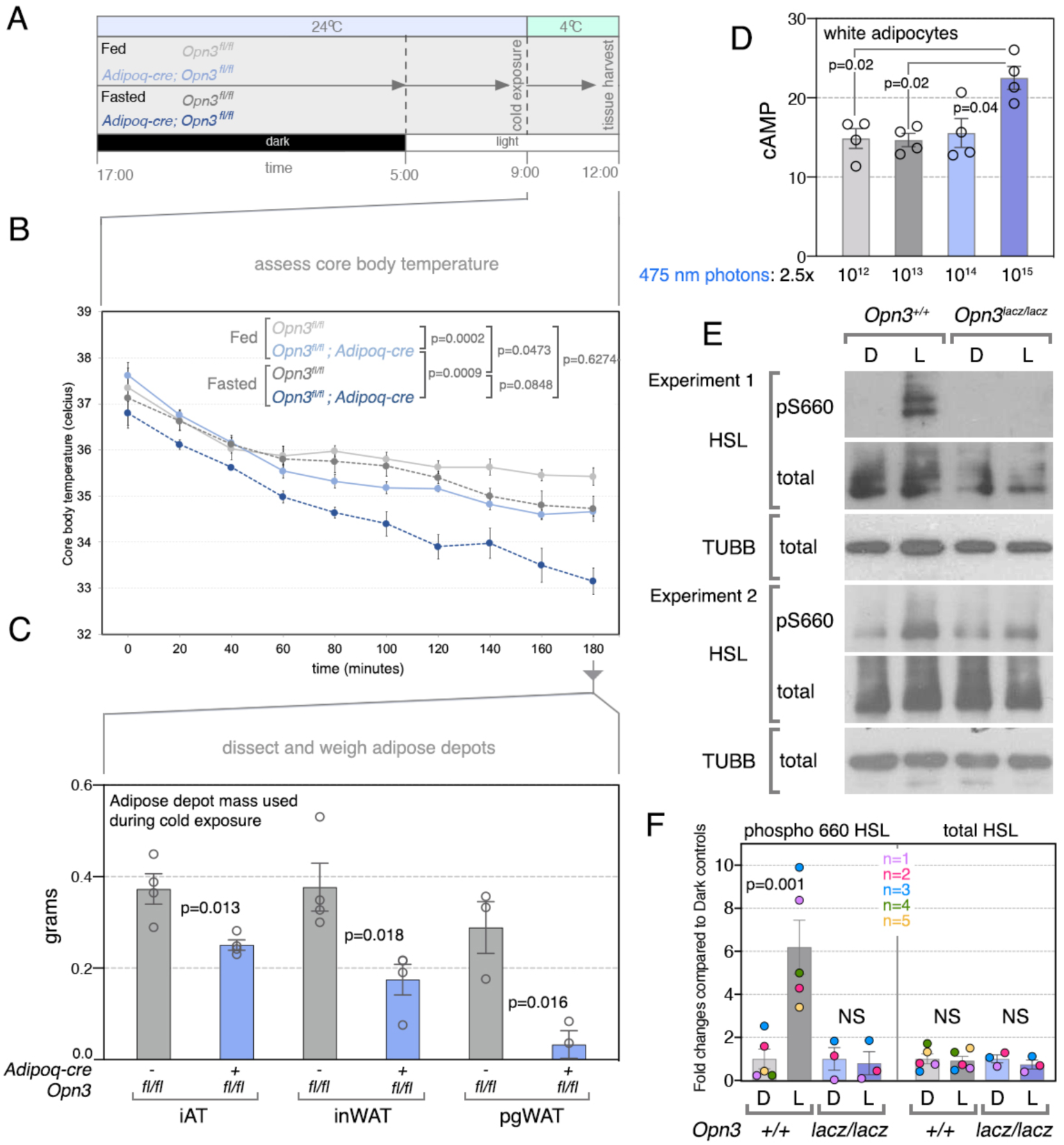
*Opn3*-dependent fat mass utilization *in vivo* and light- and *Opn3*-dependent lipolysis activation *ex vivo*. (A) Schematic describing the timeline of a fasting-cold exposure experiment that prompts mice to use fat mass for energy generation. (B) Core body temperature of fed and fasted *Opn3*^*fl/fl*^ and *Adipoq-cre; Opn3*^*fl/fl*^ mice during a 180-minute cold exposure. (C) Chart showing the fat mass used by *Opn3*^*fl/fl*^ and *Adipoq-cre; Opn3*^*fl/fl*^, fed and fasted, cold exposed mice. Compared with the controls, *Opn3* conditional null mice use less mass of iAT, inWAT and pgWAT. (D) Chart showing that cultured primary adipocytes show 475 nm light-dependent, dose response elevation of cAMP. (E) Two examples of immunoblots showing light-dependent and *Opn3*-dependent induction of phospho-660-hormone sensitive lipase (HSL) in cultured primary white adipocytes. Each set of immunblots (Experiment 1, Experiment 2) was performed using white adipocytes isolated from separate mice. (F) Quantification of phospho-HSL induction in wild type control (*Opn3*^*+/+*^, n=5 mice) and *Opn3* loss-of-function (*Opn3*^*lacz/lacz*^, n=3 mice) white adipocytes.

To test this directly, at the termination of the experiment, fat depots were dissected and weighed (Fig. 6C). On average under these conditions, cohorts of fasted control mice used 373 mg of their iAT, 377 mg of their inWAT and 289 mg of their perigonadal (pg) fat. By contrast, mice with a conditional deletion of *Opn3* in adipocytes used significantly lower quantities of fat (251 mg for iAT, 175 mg of inWAT and 33 mg of pgWAT). This indicates that adipocyte *Opn3* is required for a full utilization of fat mass. Fat mass changes under conditions of fed versus fasting state have been documented previously (Syamsunarno et al., 2014) and illustrate the need for lipolysis under conditions of nutritional deprivation.

In the second experiment, we used *ex vivo* differentiated white adipocytes from the inWAT of *Opn3*^*+/+*^ and null *Opn3*^*lacz/lacz*^ mice and assessed lipolysis pathway activation in response to stimulation with 480 nm blue photons. We chose a photon flux of 5×10^14^ photons cm^−2^sec^−1^ because this is a physiologically relevant level measured within subcutaneous adipose tissue in a mouse exposed to 1% of clear day sunlight (Fig. 2). Cultured cells of both genotypes were dark-adapted overnight and then half the samples of each genotype received 480 nm blue light stimulation for 30 minutes prior to the generation of lysates. This experimental design assesses whether white adipocytes can respond directly to light to modulate lipolysis and, if they can, whether that response is OPN3-dependent.

Conventionally, lipolysis is initiated by β-adrenergic receptor activation of Gαs and adenyl cyclase (Carey, 1998). The resulting rise in cAMP activates protein kinase A (PKA) which then phosphorylates a series of substrates including hormone sensitive lipase (HSL), perilipin (PLIN) and cAMP response element-binding protein (CREB)(Rogne and Taskén, 2014). These changes provide a signature for lipolysis pathway activation in immunoblots from cultured adipocytes. In a first experiment, we exposed primary white adipocytes isolated from inWAT to different levels of blue light stimulation and showed a dose-dependent increase in cAMP (Fig. 6D). In further analysis, we cultured primary white adipocytes from the inWAT of *Opn3*^*+/+*^ and *Opn3*^*lacz/lacz*^ mice, exposed them to either darkness or to 480 nm light, then performed immunoblotting for lipolysis pathway substrates. In adipocytes from several isolates (representing individual mice), we observed robust 480 nm light-dependent and *Opn3*-dependent phosphorylation of multiple PKA substrates including pan-phospho-PKA substrate, phospho-perilipin, phospho-HSL, and phospho-CREB (Fig. S4C). However, we also observed considerable variability in this response, perhaps because the proportion of mature adipocytes can vary. The exception was that in all culture isolates, the activation of HSL by phosphorylation at serine 660 was 480 nm light stimulated and dependent on *Opn3* (Fig. 6E). This response could be quantified: In *Opn3* wild type adipocytes, levels of light-induced phospho-HSL phosphorylation were significantly higher (Fig. 6F, gray bars) but any significant change was lost in *Opn3* null adipocytes (Fig. 6F, blue bars).

It has previously been suggested that melanopsin (OPN4) can mediate light responses in cultured primary adipocytes (Ondrusova et al., 2017). This suggested the interesting possibility that OPN3 and OPN4, both blue light responsive, might act in concert to regulate white adipocyte lipolysis. To explore this, we assessed *Opn4* expression using *Opn4*^*cre*^ (Ecker et al., 2010) to activate the *Ai14* cre reporter. Cryosections of inWAT and iBAT did not show any indication of lineage marking in adipocytes (Fig. S4E, F) even though the retina, an expected site of *Opn4* expression (Berson et al., 2002; Hattar et al., 2002) showed robust signal (Fig. S4D). To assess the possibility that melanopsin might be involved in the thermogenesis response, we measured core body temperature in cold-exposed cohorts of *Opn4* wild type and null mice. This did not reveal any significant differences (Fig. S4G), an outcome indicating that in the context of this assay, OPN4 is not essential for acute light-dependent responses in the regulation of body temperature. Finally, we also isolated primary white adipocytes from control and *Opn4* null mice, performed a blue light stimulation and showed a robust induction of phospho-HSL in both genotypes (Fig. S4H). These data are inconsistent with an *in vivo* role for melanopsin in local light responses in adipocytes in mice.

Combined, the analysis we describe identifies encephalopsin/OPN3 as a key adipocyte light sensor that regulates that activation of HSL, an enzyme central to the lipolysis response. We propose that this activity explains adipose tissue changes and the adaptive thermogenesis deficit characteristic of the *Opn3* mutant mice.

## Discussion

The analysis we present has assessed the function of Opsin 3 (Blackshaw and Snyder, 1999; Koyanagi et al., 2013b)(encephalopsin) in the mouse. *Opn3* is expressed is a wide variety of tissue types (Blackshaw and Snyder, 1999; Nissila et al., 2012; Regard et al., 2008; Sikka et al., 2016), but here we show that OPN3 mediates photic regulation of adipocyte function. Extraocular photoreception has been described in many species including vertebrates such as fish (Kojima and Fukada, 1999; Sato et al., 2016) and birds (Nakane et al., 2010). To date, however, there are only a few suggestions of extraocular light reception in mammals. Notably, it has been shown that non-visual opsins function within cultured adipocytes (Ondrusova et al., 2017), and within cultured skin cells (Regazzetti et al., 2018) and can also mediate a vasorelaxation response (Sikka et al., 2014, 2016).

### OPN3 activity mediates a light-dependent pathway that regulates energy metabolism

According to several lines of evidence a major phenotype of *Opn3* germ line null mice is deregulation of lipid homeostasis. This is evident in the transcriptome analysis from WAT where changes cluster within the PPAR pathway and the electron transport chain. Furthermore, *Opn3* germ line null mice show an adipose tissue phenotype that in WAT includes increased cell size, low brite adipocyte content, low NAD and low UCP1, and in iBAT comprises ETC complex perturbations and low UCP1. Notably, *Opn3* germ line null mice show a thermogenesis deficit and become hypothermic upon cold exposure.

OPN3 has all the crucial molecular characteristics of the opsin family of light-responsive G-protein coupled receptors (Buscone et al., 2017; Koyanagi et al., 2013b). Thus, one hypothesis to explain the *Opn3* phenotype was that mice normally use OPN3 as the detector in a pathway that decodes light information to regulate energy homeostasis. As a first step in testing this hypothesis, we simply raised C57Bl/6J mice in “minus blue” conditions that exclude the 480 nm light known to stimulate OPN3 (Koyanagi et al., 2013b). Remarkably, “minus blue” reared mice show the same abnormal WAT histology and low NAD, as well as reduced BAT ETC complexes, low UCP1 and thermogenesis deficit characteristic of the *Opn3* germ line null. This outcome is consistent with the existence of a novel OPN3-dependent, light decoding metabolic regulation pathway.

A characteristic of non-canonical opsins is that they can mediate acute responses to light. For example, melanopsin (OPN4) mediates the pupillary light reflex (Hattar et al., 2002) and light aversive behavior in neonatal mice (Johnson et al., 2010). To determine whether OPN3 could mediate light responses over a similar timescale we determined whether the thermogenesis response could be influenced by acute light changes. When we withdrew blue light during cold exposure, we showed that wild type mice rapidly reduced their body temperature to the abnormally low body temperature of *Opn3* null mice. In this assessment, the body temperatures of *Opn3* null and wild type mice were indistinguishable in “minus blue” and this indicated that OPN3 activity fully accounted for the acute enhancement of body temperature by blue light.

It has been noted that acute light stimulation enhances body temperature in humans (Cajochen et al., 2000; Dijk et al., 1991) and that this response is mediated by 460 nm and not 550 nm light (Cajochen et al., 2005). Though it has reasonably been assumed (because circadian pathways also regulate body temperature) that this reflects the activity of melanopsin (Cajochen et al., 2005) the current analysis suggests the alternative hypothesis that OPN3-dependent light responses are important to this physiology. Experiments in *Drosophila* demonstrate that acute light exposure causes elevated temperature preference (Head et al., 2015) suggesting that this configuration of light information decoding is deeply conserved. Finally, it is very likely that the activity of OPN3 in the light-dependent regulation of metabolic pathways and body temperature will be tightly integrated with circadian systems that are dependent on OPN4 and the ocular photic input pathways that also regulate this physiology.

### White adipocytes are a crucial site of OPN3 activity

Many different cell types express *Opn3* (Blackshaw and Snyder, 1999; Nissila et al., 2012; Regard et al., 2008; Sikka et al., 2016) and this raises the possibility that extraocular light reception in mammals is commonplace. In the current study we have shown, using light response assays in conditional deletion mice and in isolated cells, that white adipocytes are a crucial site of OPN3 function. We show that mice with an adipocyte conditional deletion of *Opn3* (*Adipoq-cre; Opn3*^*fl/fl*^) phenocopy the germ line null with WAT histological features and the thermogenesis response deficit. This includes the demonstration that adipocyte conditional *Opn3* null mice are hypothermic upon cold exposure and, unlike control mice, do not change their body temperature in response to acute withdrawal of blue light. Conditional deletion of *Opn3* in brown adipocytes (using *Ucp1-cre*) did not produce a thermogenesis consequence indicating that *Opn3* activity in this cell type was not required. These findings raised the question of which OPN3-dependent white adipocyte pathway is required for a normal thermogenesis response.

Prompted in part by transcriptome analysis, we showed that lipolysis in cultured white adipocytes is enhanced by blue light in an OPN3-dependent manner. The lipolysis pathway is conventionally initiated by β-adrenergic receptor and Gαs activation of cyclic AMP (Carey, 1998). In turn, cAMP activates PKA and this results in a series of substrate phosphorylations that activate HSL and perilipin, activities central to the mobilization of fatty acids and their use as fuel. As illustrated by the lower than normal body temperature that results when lipid mobilization enzymes are compromised (Schreiber et al., 2017), lipolysis is an essential component of a normal thermogenesis response in mice. Blue light stimulated white adipocytes show elevated cAMP and importantly, dramatic elevation of the active, phosphorylated form of HSL, the rate limiting enzyme in the lipolysis pathway (Langin et al., 1996). This increase in phospho-HSL was *Opn3*-dependent as the response was lost in *Opn3* null adipocytes. Since lipolysis is essential for normal body temperature maintenance (Shin et al., 2017, 2018) these findings provide, at least in part, a mechanistic explanation for the OPN3-dependent thermogenesis deficit. The reduced ability of adipocyte conditional *Opn3* null mice to use fat mass in response to fasting and cold exposure is consistent with a role for OPN3 in enhancing lipolysis *in vivo*. In mice, a nocturnal species, plasma FFA and glycerol are elevated during the day, the animal’s rest phase (Shostak et al., 2013b, 2013a). Elevated lipolysis during the light phase is likely an adaptation for providing energy when mice are inactive and not consuming food as an energy source.

Both the primary amino acid sequence and expression pattern of OPN3 are highly conserved in mammals. If the light-OPN3 adipocyte pathway exists in humans, there are potentially broad implications for human health. Our modern lifestyle subjects us to unnatural lighting spectra, exposure to light at night, shift work and jet-lag, all of which result in metabolic disruption (Fonken and Nelson, 2014; Fonken et al., 2013; Laermans and Depoortere, 2016; Opperhuizen et al., 2017). Based on the current findings, it is possible that insufficient stimulation of the light-OPN3 adipocyte pathway is part of an explanation for the prevalence of metabolic deregulation in the industrialized nations where unnatural lighting has become the norm.

## Acknowledgements

We thank Paul Speeg for excellent mouse colony management. This work was supported by NIH R01 GM124246 (EDB), NIH R01EY026921 (RVG), NIH P30EY001730 to University of Washington, the Mark J. Daily, MD Research Fund to University of Washington, and unrestricted grants to the University of Washington and from Research to Prevent Blindness. This work was also supported by grants from the NIH including R01 DK107530 (TN) R01 EY027077 (RAL, SR), R01 EY027711 (Mike Iuvone and RAL), as well as a Packard Foundation Fellowship (AMS), an American Heart Association grant (18CDA34080527)(JSG) and by funds from the Goldman Chair of the Abrahamson Pediatric Eye Institute at Cincinnati Children’s Hospital Medical Center.

## Author contributions

GN, SV, KXZ, Experimental design and analysis, manuscript preparation. YO, EDB, AH-J, ANS, BAU, JJZ, ND, SD’S, MTN, SAG, GW, RS, XM, SR, Experimental execution and analysis. NTP, MB: Electronic device design and construction. JBH: Supervision of bioinformatics analysis. TN, AS, RNVG, JS-G, RAL: Project leadership, experimental design and manuscript preparation.

## Experimental Procedures

### Mice

Animals were housed in a pathogen-free vivarium in accordance with institutional policies. Genetically modified mice used in this study were: *B6;FVB-Tg(Adipoq-cre)1Evdr/J* (Eguchi et al., 2011b)(Jax stock #010803), *Ai14* (Madisen et al., 2010)(Jax stock #007914), *Opn4* (Panda et al., 2003), and *Tg(Opn3-EGFP)JY3Gsat* (MMRRC stock number 030727-UCD). The *Ucp1*^*cre*^ mouse line used in the thermoregulation assay studies was obtained from Jackson Laboratories: *B6*.*FVB-Tg(Ucp1-cre)1Evdr/J* (Jax stock #024670). The *Opn4*^*cre*^ mouse line was obtained from the MMRRC repository as cryopreserved sperm: *Tg(Opn4-cre)SA9Gsat/Mmucd* (MMRRC stock 036544-UCD). *Opn3*^*tm2a(EUCOMM)Wtsi*^ mice were generated from C57BL/6N ES cells obtained from EUCOMM (ES clone ID: EPD0197_3_E01). The ES cells harbor a genetic modification wherein the lacz-Neomycin cassette is flanked by FRT sites and a *loxp* site separates lacz from the neomycin coding region. *Loxp* sites also flank exon 2 of *Opn3* allowing multiple mouse lines that can serve as reporter nulls, conditional floxed and null mice. The *Opn3*^*Lacz*^ reporter-null line was created by crossing *Opn3*^*tm2a(EUCOMM)Wtsi*^ mice to *FVB/N-Tg(EIIa-cre)C5379Lmgd/J* mice (Lakso et al., 1996)(Jax stock #003314). *Opn3*^*fl/fl*^ line was created by crossing the *Opn3*^*tm2a(EUCOMM)Wtsi*^ mice to *129S4/SvJaeSor-Gt(ROSA)26Sor*^*tm1(FLP1)Dym*^*/J*(Henrich et al., 2000)(Jax stock #003946) to remove the *lacZ* cassette. This means that *Opn3*^*lacz*^ mice are of mixed C57Bl6/6N, FVB/N background and that *Adipoq-cre; Opn3*^*fl*^ mice are of mixed C57Bl6/6N, 129S4/Sv, B6;FVB background. Littermate control animals were used for all experiments with the exception of C57BL/6J mice reared under different lighting conditions.

The *Opn3*^*cre*^ allele was generated in-house using CRISPR-Cas9 technology. Four gRNAs that target exon 2 of *Opn3* were selected to knock in the Cre cassette. Plasmids containing the gRNA sequence were transfected into MK4 cells (an in-house mouse cell line representing induced metanephric mesenchyme undergoing epithelial conversion). The editing efficiency of gRNA was determined by T7E1 assay of PCR products of the target region amplified from genomic DNA of transfected MK4 cells. The sequence of the gRNA that was subsequently used for the transfection is TACCGTGGACTGGAGATCCA. Sanger sequencing was performed to validate the knock-in sequence of founder mice.

The *Opn3Δ*^*Ex2*^ allele was generated in-house using CRISPR-Cas9 technology as above. Four gRNAs that target exon 2 of *Opn3* were selected. The sequences of the gRNAs are: for the 5’ end: TAGCAACGAATGCAAAGGTA GGG and ATCCACATGTTCTGCCCAGGAGG. For the 3’ end: GCGCTATGTTGGTAAGGTGT GGG and TGTGGTTTTAATCAGCACAGGGG. Out of the 6 pups derived from one injection, the founder animal had a 2203 bp deletion that was also confirmed by Sanger sequencing. The proximal breakpoint of this deletion is intron 1 (bp 175,667,424) to intron 2 (bp175,665,054) thus deleting the entirety of exon 2.

Mice were placed on normal chow diet (NCD: 29% Protein, 13% Fat and 58% Carbohydrate kcal; LAB Diet #5010) ad libitum with free access to water. For fasting experiments, mice aged 10 – 12 weeks were fasted overnight from ZT12 (lights off) to ZT3 (3 hours after lights on) for a total of 15 hours.

### Genotyping

Primer sequences and pairs for genotyping each of the alleles in this study are listed in the table below:

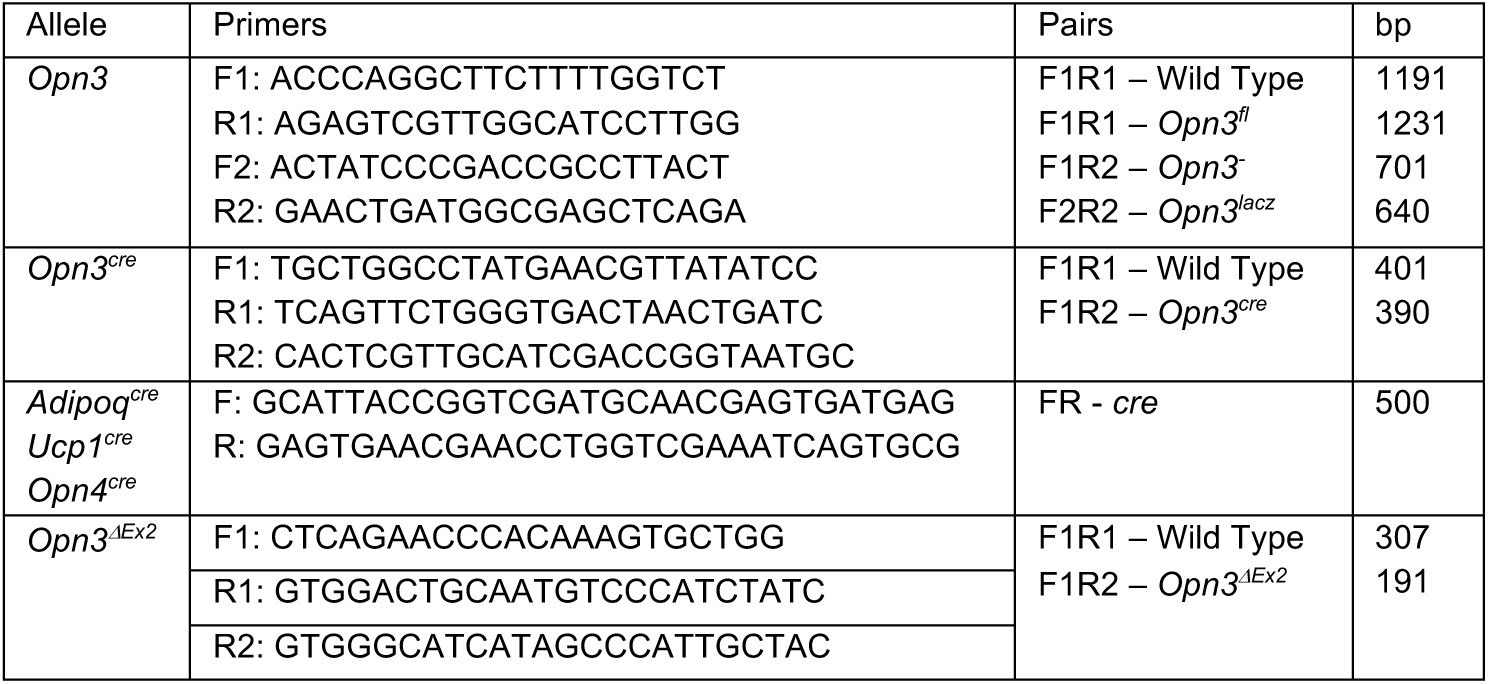

### Lighting conditions

Animals were housed in standard fluorescent lighting (photon flux 1.62×10^15^ photons cm^−2^sec^−1^) on a 12L:12D cycle except where noted. For full spectrum lighting, LEDs were used to yield a comparable total photon flux of 1.68×10^15^ photons cm^−2^sec^−1^. Spectral and photon flux information for LED lighting: near-violet (λ_max_=380 nm, 4.23×10^13^ photons cm^−2^sec^−1^ in 370-400 nm range), blue (λ_max_=480 nm, 5.36×10^14^ photons cm^−2^sec^−1^ in 430-530 nm range), green (λ_max_=530 nm, 5.82×10^14^ photons cm^−2^sec^−1^ in 480-600 nm range) and red (λ_max_=630 nm, 1.93×10^14^ photons cm^−2^sec^−1^ in 590-660 nm range). For wavelength restricted growth assessment, C57BL/6J animals were housed in 12L:12D cycle starting late gestation (embryonic day E16) either with blue (480 nm and 380 nm LEDs) or without blue (480 nm LEDs) lighting.

### Intra-adipose tissue radiometry

Fabrication of the Holt-Sweeney microprobe (HSM) is described as follows (Holt et al., 2014). The termination of one end of a 100 μm silica core fiber optic patch cable (Ocean Optics, Dunedin, FL, USA) was removed. The fiber’s furcation tubing and jacketing was stripped, and the fiber’s polyimide buffer was removed 5 cm from the fiber’s end using a butane torch. A 10 g weight was attached to the end of the fiber and then pulled upon heating with the butane torch, narrowing the diameter. The narrowed region of the fiber was then cut using carborundum paper, to yield a flat fiber end with a diameter of 30 – 50 μm. The sides of the narrowed fiber were painted with a film opaquing pen to prevent stray light from entering, while leaving a small transparent opening at the fiber tip. For structural support, this bare, tapered fiber was then secured in the tip of a pulled glass Pasteur pipette using a drop of cyanoacrylate glue, leaving only 6-9 mm of bare optical fiber protruding. A small light-scattering ball was added to the end of the tapered optical fiber for spectral scalar irradiance measurements. To do this, titanium dioxide was thoroughly mixed with a high-viscosity UV-curable resin, DELO-PHOTOBOND, GB368 (DELO Industrie Klebstoffe, Windach, Germany). The tip of a pulled fiber was quickly inserted and removed from a droplet of the resin and titanium dioxide mixture, resulting in a sphere with a diameter of approximately twice that of the tapered fiber. As all measurements from a given probe were normalized to the signal from the same probe in a gelatin blank, small variations in the probe diameter have no effect on our results. The sphere was cured for 12 h using a Thorlabs fiber coupled LED light source (M375F2, Thorlabs Inc, Newton, NJ, USA).

For intra-tissue radiometric measurements in mice, animals were anesthetized under ventilated isoflurane and placed in a mouse stereotaxic frame (Stoelting Co, Wood Dale, IL, USA). The hair overlying the intrascapular region was shaved and a small 10 mm rostrocaudal incision was made through the dorsal skin to expose the underlying tissue. A 21-gauge needle attached to the stereotaxic frame was first lowered through the intrascapular region to produce a pilot hole through the adipose tissue. Following, the Holt-Sweeney microprobe was affixed to the stereotaxic frame and lowered through the pilot hole. After the probe is in position, the dorsal skin was repositioned to cover as much of the incision site as possible without obstructing the probe’s descent. For broadband light illumination, a Thorlabs plasma light source (HPLS345, Thorlabs Inc, Newton, NJ, USA) was positioned above and in front of the mouse stereotaxic frame. The light was delivered to the animal via a 5 mm liquid light guide connected to a 2 in. collimating lens secured in a vice. The distance from the collimating lens to the animal was approximately 2 ft. Scalar irradiance measurements as a function of wavelength were obtained at the surface of the adipose tissue and at probe depth increments of 0.5 mm up to 2.5 mm. Spectral irradiance data was collected using an Ocean Optics 200-850 nm spectrometer (JAZ series, Ocean Optics, Dunedin, FL, USA).

### Immunohistochemistry and tissue processing

Animals were anesthetized under isoflurane and sacrificed by cervical dislocation. Adipose tissue depots (interscapular adipose tissue complex and inguinal WAT) were harvested and fixed in ice cold 10 % zinc formalin for 1 hour at 4°C. After washing in PBS, adipose tissue samples were prepared for cryosectioning as described previously (Berry et al., 2014). Gelatin-embedded tissues were sectioned at 16 μm in a cryostat and labeled with primary antibodies as previously described(Berry et al., 2014). Rabbit antibodies to GFP (ab13970, 1 in 500), COX4 (Genetex, GTX114330, 1 in 400, cold acetone:methanol, 1:1, post-fixed, 10 min), and UCP1 (ab10983, 1 in 500), were purchased from Abcam. Alexa 647 conjugated isolectin (1 in 300) and Alexa 546 conjugated F-actin were purchased from Invitrogen. Alexa 488 conjugated secondary antibodies (1 in 300) were purchased from Jackson ImmunoResearch.

### X-Gal staining

For X-Gal labeling, tissue samples were fixed in X-Gal fixative (1% formaldehyde, 0.2% glutaraldehyde, 2 mM MgCl_2_, 5 mM EGTA, and 0.01% Nonidet P-40) for two hours at room temperature. Tissues were cryosectioned as described above and then labeled with X-Gal. The reaction was monitored closely and stopped when background started to appear in control (wild-type) tissues. Following two washes in PBS, cryosections were imaged using a bright field microscope.

### Hematoxylin labeling and cell-size quantification

Gelatin-embedded frozen sections of inguinal WAT (as described above) were stained with hematoxylin and imaged under bright-field. Samples were imaged with a rhodamine filter to assess adipocyte size distribution. Using the free hand selection tool on ImageJ, adipocytes were outlined and the area measured in μm^2^. Cell size distribution was determined by quantifying 60 cells from at least 10 regions, for a total of approximately 600 cells per animal. Cell areas were binned into 200 μm^2^ intervals and the frequency of total cells (%) charted for each interval.

### Pre-Adipocyte differentiation and light induction

IngWAT dissociation and extraction of stromal vascular fraction was performed as described before (Liu et al., 2017). Briefly, the inguinal fat pads were collected in PBS and digested in 1.5 mg/ml Collagenase A in PBS with 4% BSA and penicillin/streptomycin at 37°C, with intermittent agitation over 40 minutes. The stromal vascular fraction was extracted by passing the enzymatically dispersed cells through a 100 μm cell strainer and cultured in basal media (DMEM containing 10% fetal bovine serum and penicillin/streptomycin). For differentiation, the stromal vascular cells were plated on day1 such that the cells reached confluency on day 3. On day 4, the basal media was replaced with induction media containing Insulin (100 nM), Rosiglitazone (1 μM), IBMX (0.5 mM) and Dexamethasone (2 μg/ml) in basal media. Thereafter, the differentiating cells were maintained in basal media containing insulin (100 nM) until the day of experimentation.

Light inductions to assay lipolysis responses were typically done on day 13 – day 15 of differentiation. For this, cultures were moved to a dark, 37°C incubator, protected from light, overnight. The next day, the cultures were serum-starved, under dim red light, where the complete basal medium was washed out using serum-free medium (at least three washes) and the cells were left in serum-free media for 3 hours, before light inductions. For light inductions, half the *Opn3*^*+/+*^ and *Opn3*^*Lacz/Lacz*^ cultures were left in the dark incubator, while the rest were moved to an adjacent incubator that housed a light set-up to deliver 5×10^14^ photons/cm^2^/sec of 480 nm wavelength. The culture conditions in the two incubators were comparable except for the lighting. Light inductions were carried out for 30 minutes, after which the cells were washed in PBS and snap frozen by immersing the culture plates in liquid nitrogen and frozen at −80°C until lysate preparations for Western blotting.

For the light-induced cyclic AMP response, *in vitro* differentiated *Opn3*^*+/+*^ and *Opn3*^*Lacz/Lacz*^ adipocytes were used between days 7 and 10 of differentiation. The cells were incubated with 9-cis-Retinal (5 μM) the day before the assay and one hour before the light induction, the cells were incubated in fresh DMEM without phenol red. The light pulses (465 nm) were delivered for 30 minutes with varying intensities as indicated in the results. The cells were then harvested to quantify cAMP levels by direct immunoassay (fluorometric kit by Abcam, ab138880) as per manufacturer’s instructions. Briefly, all samples and standards (50 ul each) were tested in duplicates, to which 25 μl of 1x HRP-cAMP was added. The plates were incubated at room temperature for 2 hours and after the washing steps, 100 μl of AbRed indicator was added. The plates were incubated for 1 hour and the fluorescence was measured at Ex/Em=540/590 nm using a Biotek Synergy4 microplate reader.

### Western Blotting

Western blots were performed using standard protocols. Adipose tissue lysates were made in NP40 lysis buffer: 150 mM NaCl, 1% NP40, 50 mM Tris 8.0 with phosphatase inhibitors. Lysates were prepared by sonication and were separated from the overlying fat layer by three rounds of centrifugation. For cultured cells, RIPA lysis buffer containing proteinase and phosphatase inhibitors was added onto frozen cells and left on ice for 30 minutes with intermittent agitation for protein extraction. The lysates were cleared of lipid using two rounds of 4°C centrifugation. After the BCA method of protein quantification, lysates were boiled in Laemmli sample buffer (4% SDS, 20% glycerol, 10% 2-mercaptoethanol, 0.004% bromophenol blue and 0.125 M Tris HCl, pH 6.8). Blots were incubated with OxPhos antibodies (Thermofisher 45-8099 1:1000), UCP-1 (Abcam ab10983), beta-tubulin (ab21057 at 1:2000) and lipolysis pathway proteins from Cell Signaling at 1:1000 -phospho 660-HSL (#4126), total HSL (#4107), phospho PKA substrate (#9624), phospho (Thr197) PKA (#5661), Total PKA (#4782), phospho CREB (#9198), phospho ERK (#9101) and phospho perilipin (Vala Sciences; 4856 at 1:500). HRP-conjugated secondary antibodies (Thermoscientific) were used at 1:5000 dilution and detected by enhanced chemiluminescence (ThermoFisher Scientific).

### Microarray analysis

Interscapular adipose tissue complex and inguinal white adipose tissue from P16 mice were harvested at one hour after lights on (ZT1) and snap frozen on dry ice. Tissue pieces were homogenized in TRIzol (TriReagent Invitrogen) using RNAse-free Zirconium oxide beads (2.0 mm) in a TissueLyser II (Qiagen). Phase separation was achieved using chloroform and RNA in the aqueous phase was precipitated using ethanol. RNA was purified by column method using GeneJET RNA purification kit (ThermoFisher Scientific #K0732) and eluted into RNAse-free water. RNA quality was assessed using the Agilent 2100 Bioanalyzer and an RNA-integrity number cut-off of 7 was applied for selecting samples for microarray assay. RNA from biological triplicates were submitted for microarray assay (ClariomD, Affymetrix) to the Technology Center for Genomics and Bioinformatics, University of California, Los Angeles.

Data analysis including normalization, gene expression changes and gene-enrichment analysis was performed using AltAnalyze, developed by Nathan Salomonis at Cincinnati Children’s Hospital Medical Center. AltAnalyze uses the robust multi-array average method of normalization. Briefly, the raw intensity values are background corrected, log2 transformed and then quantile normalized. Next, a linear model is fit to the normalized data to obtain an expression measure for each probe set on each array. Gene expression changes greater than 1.1 fold were calculated using unpaired t-test, where a p-value <0.05 was used as a cut-off.

### Quantitative RT-PCR

Intrascapular adipose depots were harvested immediately following cold challenge assays. Snap frozen tissue was homogenized and processed for RNA as described above. RNA was treated with RNase-free DNase I (ThermoFisher Scientific #EN0521) and cDNA was synthesized using a Verso cDNA synthesis kit (ThermoFisher Scientific AB1453/B). Quantitative RT-PCR was performed with Radiant™ SYBR Green Lo-ROX qPCR mix (Alkali Scientific Inc.) in a ThermoFisher QuantStudio 6 Flex Real-Time PCR system. Primer information for quantitative PCR is included in the Table. Relative expression was calculated by the ΔΔCT method using *Tbp* (TATA binding protein) as the normalizing gene. Statistical significance was calculated by an unpaired t-test, using a p-value cutoff of <0.05.

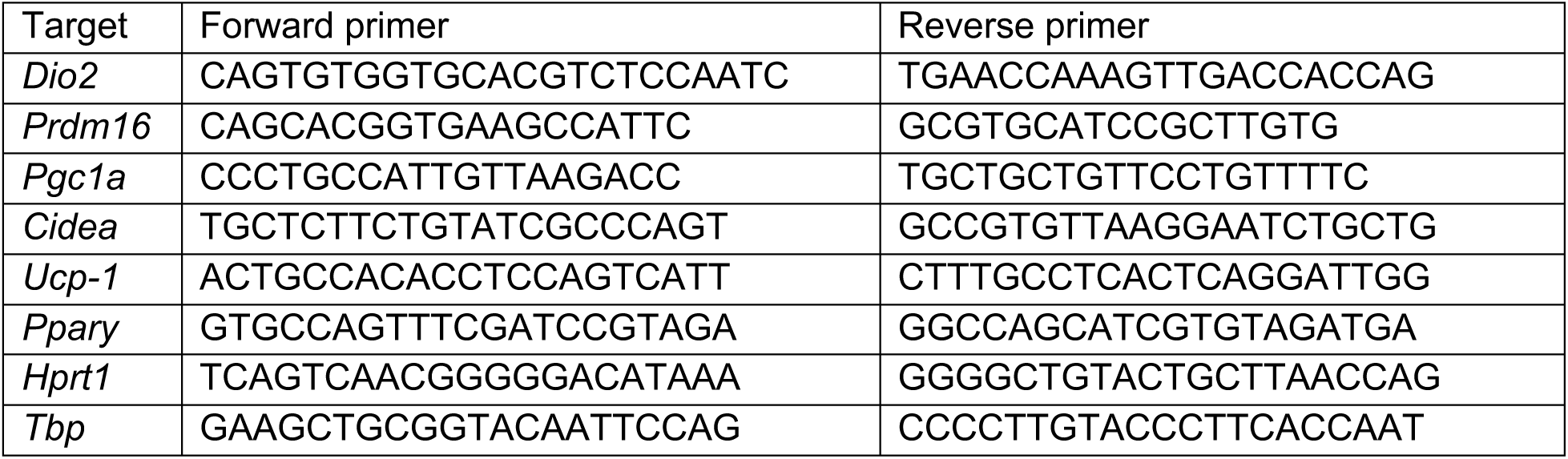

### Transmission Electron Microscopy

Freshly dissected adipose tissues were collected and 1 mm samples from approximately similar areas were fixed in 2% glutaraldehyde, 1% paraformaldehyde in PBS for 1 hour at room temperature before processing and sectioning for transmission electron microscopy as described before (Cinti S, Zingaretti MC, Cancello R, Ceresi E, 2001).

### NAD/NADH Quantification

NAD levels were measured using NAD/NADH assay kit from Abcam (ab65348). Briefly, tissues samples (inguinal adipose tissue and liver) from P16 mouse pups were snap frozen in liquid nitrogen, homogenized in NADH/NAD extraction buffer and filtered through a 10kD spin column (ab93349) to remove enzymes. Assay procedure was followed per kit instructions and levels of NADH and NAD+ were determined normalized to tissue weight.

### Thermoregulation assay

Core body temperature assessment upon acute cold exposure was performed on control and experimental mice with the *Opn3* reporter null (*Opn3*^*+/+*^ and *Opn3*^*lacz/lacz*^), with the exon 2 deletion on the C57BL/6J background (*Opn3*^*ΔEx2*^), with pan-adipocyte conditional deletion of *Opn3* (*Opn3*^*fl/fl*^ and *Adipoqcre; Opn3*^*fl/fl*^), and with brown adipocyte conditional deletion of *Opn3* (*Opn3*^*fl/fl*^ and *Ucp1cre; Opn3*^*fl/fl*^). In addition, C57BL/6J mice reared under wavelength restriction (with or without blue, as described previously) were subject to this assay. Littermates were separated from their home cage and individually housed in a home-built lighting chamber situated in an electronically monitored 4°C cold room for 3 or 5 hours depending on the assay. While the mouse was conscious, body temperature was measured rectally with a RET-3 Microprobe Thermometer (Kent Scientific) every 20 minutes for the duration of the assay. Food and water were available *ad libitum* for all mice except when *Adipoqcre; Opn3*^*fl/fl*^ mice were fasted overnight, where food withdrawal was maintained during the cold assay. The thermo probe operator was blinded to mouse genotype and prior temperature measurements throughout the study. At the end of the cold exposure, mice were euthanized and relevant tissues were collected. The 3-hour cold exposure assays subjected mice to either a red (630 nm) and violet (380 nm) LED illumination combination (RV), or a red (630 nm), blue (480 nm) and violet (380 nm) LED combination (RBV). For the 5-hour cold exposure assays, the entirety of the 3-hour assay was extended by 2 hours following withdrawal of the 480 nm wavelength LED illumination. Two different ages of animals, postnatal day 21 and 6 week adults, were selected for these cold exposure assays. The order of cage placement was randomized at this time, such that the thermo probe operator remained blinded. For all cold exposure assays involving fed or overnight fasted *Adipoqcre; Opn3*^*fl/fl*^ animals, intrascapular (iAT), inguinal (inWAT), and perigonadal (pgWAT) adipose tissues were harvested. Following animal euthanasia, the fat pads were manually dissected, and their weight recorded. For fat depots with left and right pads, both were harvested and weighed, and the average recorded per animal.

### Data analysis

Statistical analyses were performed using GraphPad Prism version 4.00 (GraphPad Software), Microsoft Excel and MATLAB (Figure 2). Rectal temperature charts (Figures 4, 5) were analyzed via one-way repeated measures ANOVA. For adipose weights and immunoblot quantification, a two-tailed distribution, two-sample unequal variance *t*-test was used to determine the statistical significance between two independent groups.

