## Supplementary Figure 1-4 for "Adaptive thermogenesis in mice requires adipocyte light-sensing via Opsin 3"

### Supplementary Figures and Legends

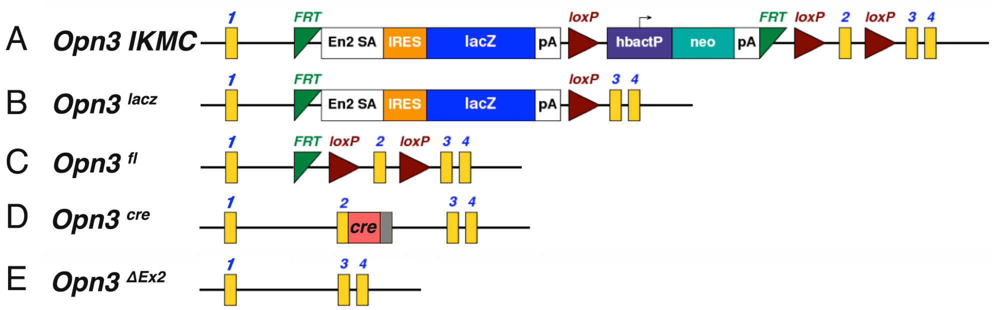


**Supplementary Figure 1. *Opn3* alleles and *Opn4* null neonatal growth**
(A-D) Schematics of the *Opn3* alleles used in this study. (A) The *Opn3* allele as targeted in ES cells by the International Knockout Mouse Consortium. Exons are numbered yellow boxes. *FRT*, FLP recombinase site-specific recombination sites. En2 SA, *Engrailed 2* splice acceptor. IRES, internal ribosome entry sequence. *Lacz*, β-galactosidase open reading frame. pA, polyadenylation signal. loxP, cre recombinase site specific recombination sequences. hbactP, human β-actin promoter. Right-facing arrow indicates the start point of transcription for hbactP. neo, the neomycin resistance gene. (B) The *Opn3*$^{lacz}$ allele generated after germ-line recombination at the *loxP* sites. (C) The *Opn3*$^{fl}$ allele generated after germ-line recombination at the *FRT* sites. (D) The *Opn3*$^{cre}$ knock-in allele was generated using CRISPR/Cas9 mediated insertion of a cre recombinase open reading frame into exon 2. (E) The *Opn3*$^{\Delta Ex2}$ allele was generated using CRISPR/Cas9 mediated deletion of exon 2.

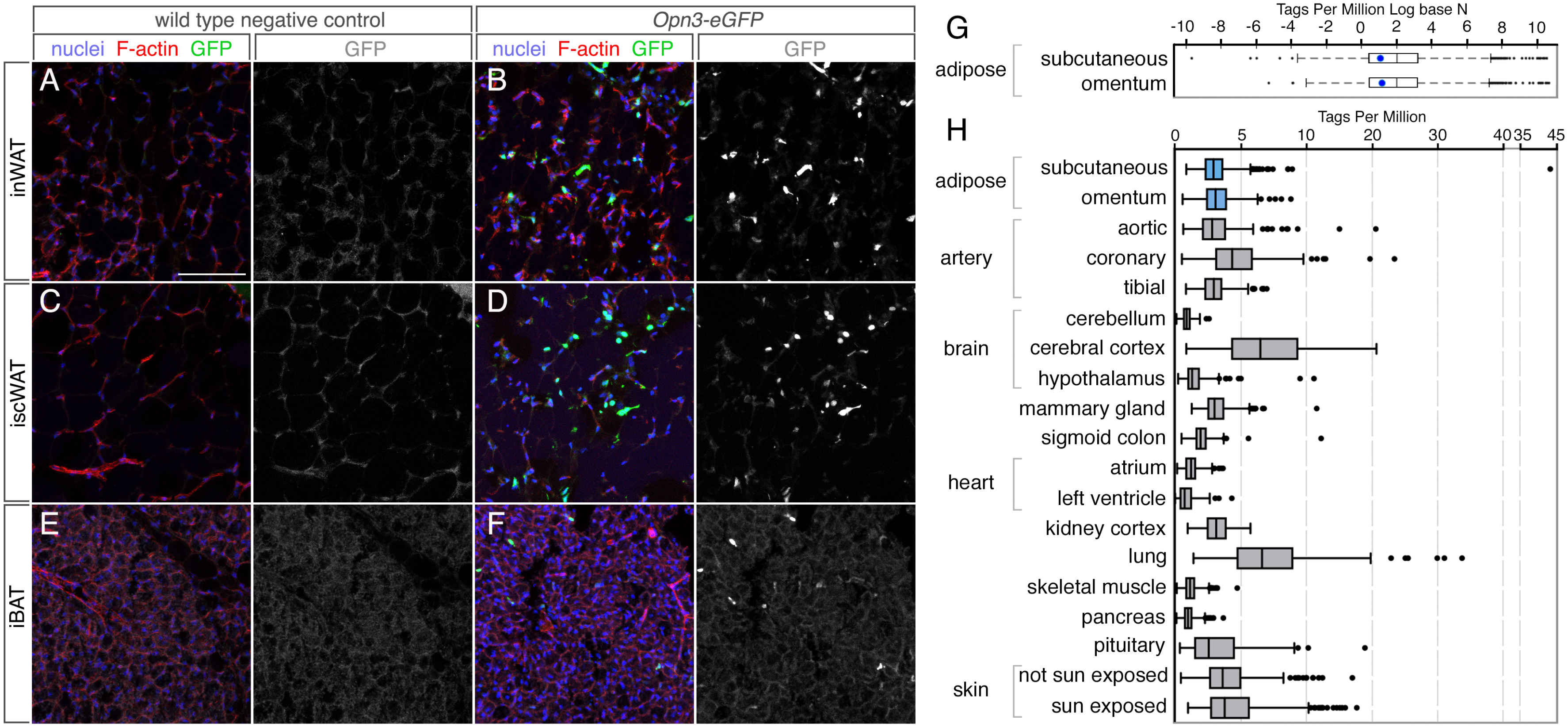


**Supplementary Figure 2. Expression of *Opn3-eGFP* reporter in adipose tissue: Expression of *OPN3* in human tissue.**

(A-F) Panels show cryosections of adipose tissue from P16 mice labeled with Hoechst33258 to detect nuclei (blue), with fluorochrome-conjugated Phalloidin to detect F-actin (red), and with anti-GFP antibodies (green, gray). (A, B) Inguinal white adipose tissue (inWAT), (C, D) interscapular white adipose tissue (iscWAT), (E, F) interscapular brown adipose tissue (iBAT). For clarity, the green channel detecting GFP is shown in grayscale. White, 100 μm scale bar in (A) applies to all immunofluorescence panels. (G) Box and whiskers plot showing Tags Per MIllion (TPM)(Log base N) sequencing reads for the *OPN3* transcript (blue dot) versus all other transcripts in human subcutaneous and omental adipose tissue. The box defines the interquartile data range (IQR, the difference between the 25$^{th}$ and 75$^{th}$ percentiles), and the line within the box, the median. Box plot whiskers are plotted according to the Tukey method. The *OPN3* transcript is detected at 2.9 TPM for subcutaneous adipose and 3.1 TPM for omental adipose. The median value for all transcripts in these adipose tissues is 7.6 TPM for subcutaneous and 7.0 for omental. (H) Box and whiskers plot showing Tags Per MIllion sequencing reads for human *OPN3* transcript in the indicated tissues. Subcutaneous and omental adipose tissue median *OPN3* expression is in the mid-range of expression values.

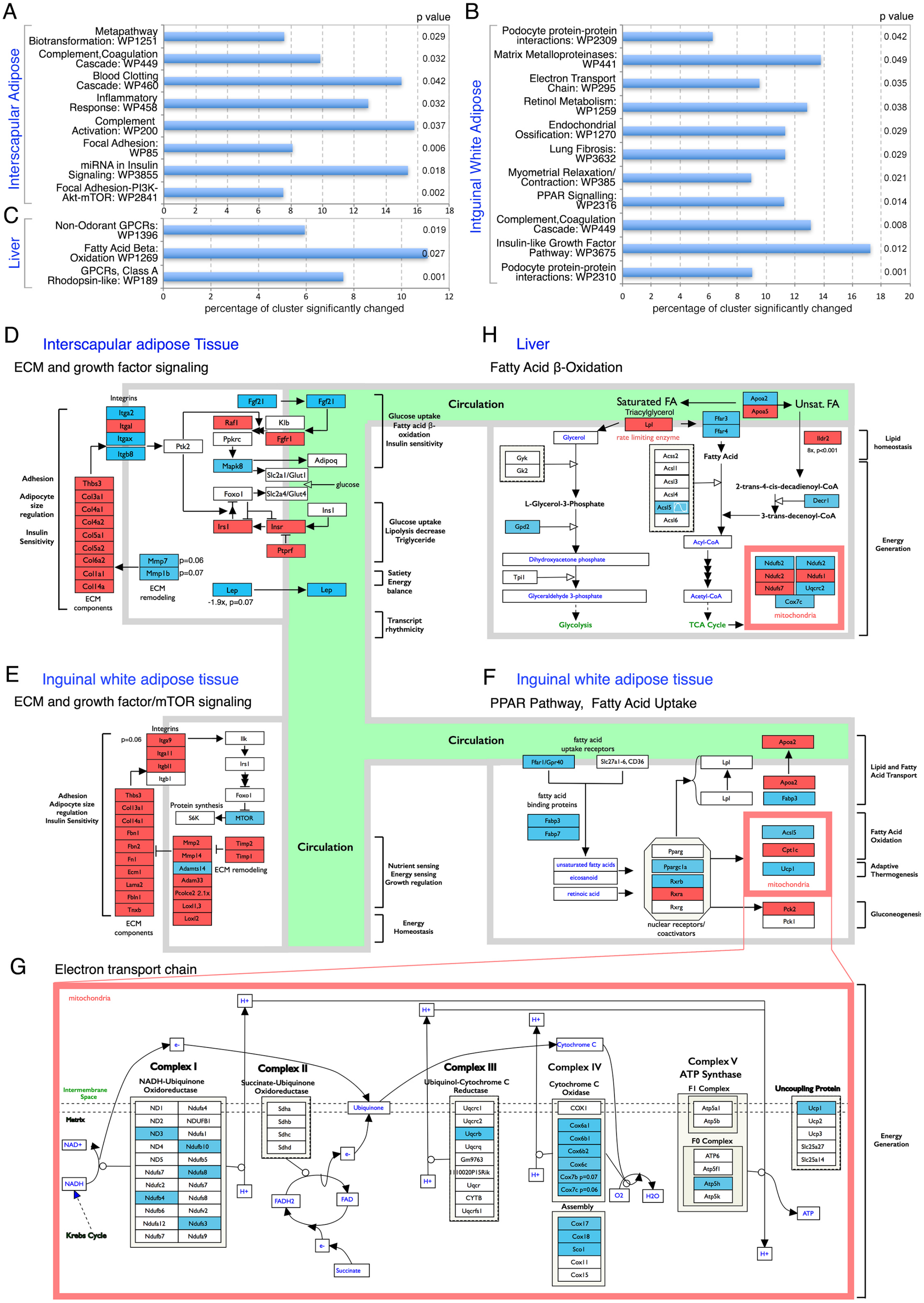

A
Interscapular Adipose
p value
Metapathway Biotransformation: WP1251
Complement,Coagulation Cascade: WP449
Blood Clotting Cascade: WP460
Inflammatory Response: WP458
Complement Activation: WP200
Focal Adhesion: WP85
miRNA in Insulin Signaling: WP3855
Focal Adhesion-PI3K-Akt-mTOR: WP2841
0.029
0.032
0.042
0.032
0.037
0.006
0.018
0.002
0 2 4 6 8 10 12 14 16 18
C
Liver
Non-Odorant GPCRs: WP1396
Fatty Acid Beta: Oxidation WP1269
GPCRs, Class A Rhodopsin-like: WP189
0.019
0.027
0.001
0 2 4 6 8 10 12
percentage of cluster significantly changed
B
Intguinal White Adipose
p value
Podocyte protein-protein interactions: WP2309
Matrix Metalloproteinases: WP441
Electron Transport Chain: WP295
Retinol Metabolism: WP1259
Endochondrial Ossification: WP1270
Lung Fibrosis: WP3632
Myometrial Relaxation/ Contraction: WP385
PPAR Signalling: WP2316
Complement,Coagulation Cascade: WP449
Insulin-like Growth Factor Pathway: WP3675
Podocyte protein-protein interactions: WP2310
0.042
0.049
0.035
0.038
0.029
0.029
0.021
0.014
0.008
0.012
0.001
0 2 4 6 8 10 12 14 16 18 20
percentage of cluster significantly changed
D
Interscapular adipose Tissue
ECM and growth factor signaling
Integrins
Itga2
Itgal
Itgax
Itgb8
Ptk2
Fgf21
Fgf21
Raf1
Klb
Ppkrc
Fgfr1
Mapk8
Adipoq
Slc2a1/Glut1
glucose
Foxo1
Slc2a4/Glut4
Ins1
Irs1
Insr
Ptprf
Adhesion
Adipocyte size regulation
Insulin Sensitivity
Thbs3
Col3a1
Col4a1
Col4a2
Col5a1
Col5a2
Col6a2
Col1a1
Col14a
ECM components
Mmp7 p=0.06
Mmp1b p=0.07
ECM remodeling
Lep
Lep
-1.9x, p=0.07
Glucose uptake
Fatty acid β-oxidation
Insulin sensitivity
Glucose uptake
Lipolysis decrease
Triglyceride
Satiety
Energy balance
Transcript rhythmicity
H
Liver
Fatty Acid β-Oxidation
Circulation
Saturated FA
Triacylglycerol
Lpl
rate limiting enzyme
Apoa2
Apoa5
Unsat. FA
Ffar3
Ffar4
Fatty Acid
Glycerol
Gyk
Gk2
Acss2
Acsl1
Acsl3
Acsl4
Acsl5
Acsl6
Ildr2
8x, p<0.001
2-trans-4-cis-decadienoyl-CoA
Decr1
3-trans-decenoyl-CoA
L-Glycerol-3-Phosphate
Gpd2
Acyl-CoA
Dihydroxyacetone phosphate
Tpi1
Glyceraldehyde 3-phosphate
Acetyl-CoA
Glycolysis
TCA Cycle
Ndufb2
Ndufs2
Ndufc2
Ndufs1
Ndufs7
Uqcrc2
Cox7c
mitochondria
Lipid homeostasis
Energy Generation
E
Inguinal white adipose tissue
ECM and growth factor/mTOR signaling
Integrins
p=0.06
Itga9
Itga11
Itgbl1
Itgb1
Ilk
Irs1
Foxo1
Protein synthesis
S6K
MTOR
Adhesion
Adipocyte size regulation
Insulin Sensitivity
Thbs3
Col13a1
Col14a1
Fbn1
Fbn2
Fn1
Ecm1
Lama2
Fbln1
Tnxb
ECM components
Mmp2
Mmp14
Adamts14
Adam33
Pcolce2 2.1x
Loxl1,3
Loxl2
Timp2
Timp1
ECM remodeling
Circulation
Nutrient sensing
Energy sensing
Growth regulation
Energy Homeostasis
F
Inguinal white adipose tissue
PPAR Pathway, Fatty Acid Uptake
Circulation
fatty acid uptake receptors
Ffar1/Gpr40
Slc27a1-6, CD36
fatty acid binding proteins
Fabp3
Fabp7
Lpl
Lpl
Apoa2
Apoa2
Fabp3
unsaturated fatty acids
eicosanoid
retinoic acid
Pparg
Ppargc1a
Rxrb
Rxra
Rxrg
nuclear receptors/ coactivators
Acsl5
Cpt1c
Ucp1
mitochondria
Pck2
Pck1
Lipid and Fatty Acid Transport
Fatty Acid Oxidation
Adaptive Thermogenesis
Gluconeogenesis
G
Electron transport chain
mitochondria
H+
H+
H+
e-
Cytochrome C
Complex I
NADH-Ubiquinone Oxidoreductase
Complex II
Succinate-Ubiquinone Oxidoreductase
Complex III
Ubiquinol-Cytochrome C Reductase
Complex IV
Cytochrome C Oxidase
Complex V
ATP Synthase
F1 Complex
Uncoupling Protein
Intermembrane Space
Matrix
Ubiquinone
NAD+
NADH
Krebs Cycle
ND1
ND2
ND3
ND4
ND5
Ndufa7
Ndufc2
Ndufb4
Ndufb6
Ndufa12
Ndufb7
Ndufa4
NDUFB1
Ndufa1
Ndufb10
Ndufb5
Ndufa8
Ndufs7
Ndufs8
Ndufv2
Ndufs3
Ndufa9
Sdha
Sdhb
Sdhc
Sdhd
e-
FADH2
FAD
e-
Succinate
Uqcrc1
Uqcrc2
Uqcrb
Uqcrq
Gm9763
1110020P15Rik
Uqcr
CYTB
Uqcrfs1
COX1
Cox6a1
Cox6b1
Cox6b2
Cox6c
Cox7b p=0.07
Cox7c p=0.06
Assembly
Cox17
Cox18
Sco1
Cox11
Cox15
O2
H2O
Atp5a1
Atp5b
F0 Complex
ATP6
Atp5f1
Atp5h
Atp5k
Ucp1
Ucp2
Ucp3
Slc25a27
Slc25a14
ATP
H+
Energy Generation

**Supplementary Figure 3. Transcriptome analysis of *Opn3* null mice**

(A-C) Charts show, for interscapular adipose tissue (A), inguinal white adipose tissue (B) and liver (C), the WikiPathways (text labels and WP numbers) in which there was significant clustering (according to Z-score) of significantly changed transcripts in *Opn3*$^{+/+}$ versus *Opn3*$^{lacz/lacz}$ tissue. The chart horizontal axis shows the percentage of significantly changed transcripts for a given pathway. Fisher's exact test p-values are listed on the right of the chart. (D-H) Panels show schematically, in the context of known pathways, *Opn3*-dependent changes in transcripts for the indicated genes in iAT (D) inWAT (E-G) and liver (H). Each box represents a transcript that is significantly ($p<0.05$) up-regulated (red) or down-regulated (blue) in the *Opn3* null. For four transcripts (*Mmp7*, *Mmp1b*, *Lep* (D) and *Itga9* (E)) where the p-value did not quite reach $p<0.05$, the p-value is listed. The circulation is modelled by the green area. All panels are modified versions of Wikipathway schematics for Focal Adhesion-PI3K-mTOR Signaling pathway (WP2841, (D, E)), for the PPAR Signaling Pathway (WP2316, (F)), for the Electron Transport Chain (WP295, (G)) and for the Fatty Acid β-Oxidation Pathway (WP1269, (H)).

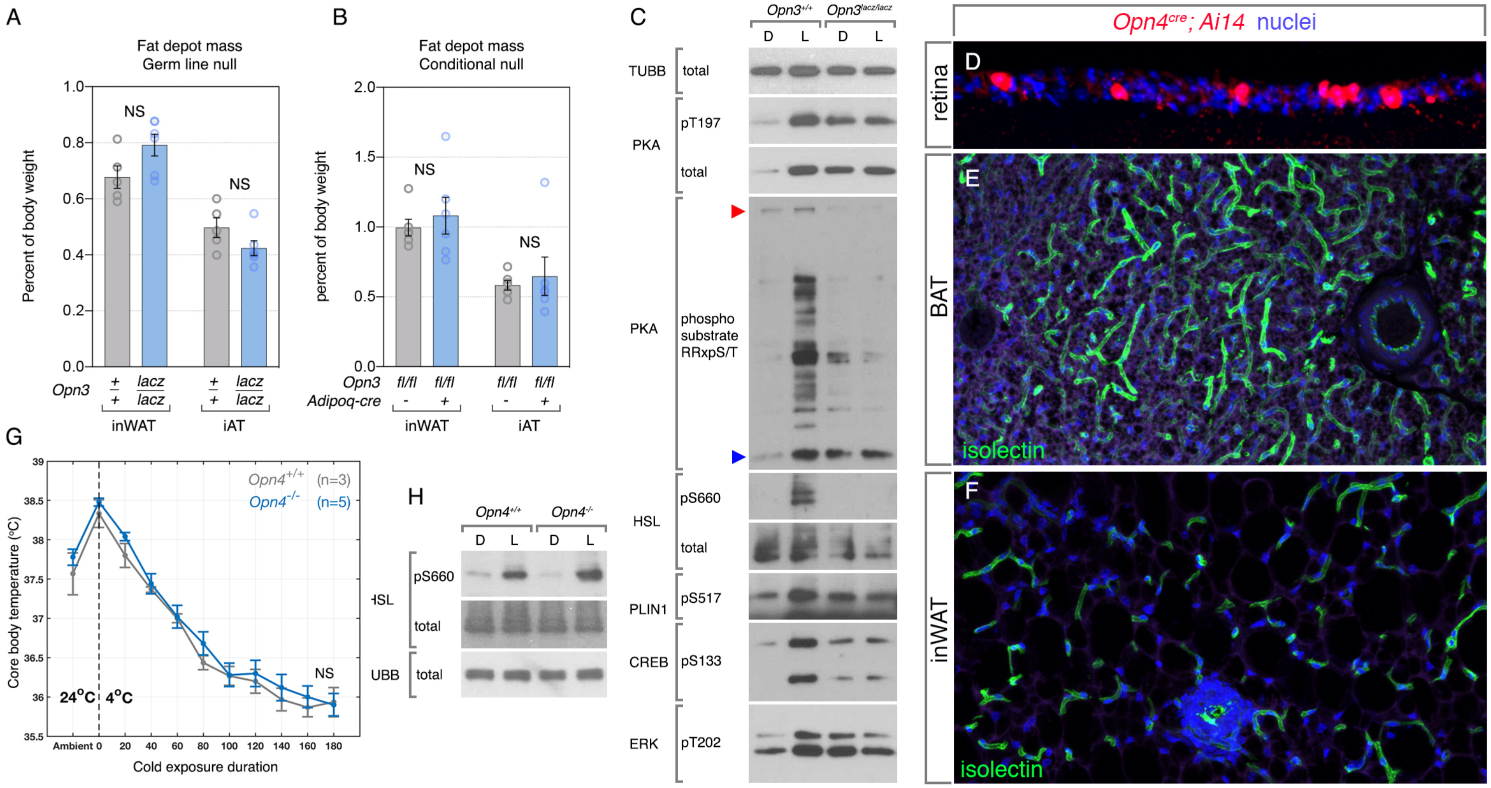


**Supplementary Figure 4: *Opn3* versus *Opn4* involvement in adipocyte responses**

(A, B) Charts showing fat mass for inWAT and iAT for control and Opn3 germ line and adipocyte conditional null mice as labeled. (C) Immunoblots showing *Opn3*- and blue light-dependent lipolysis activation in cultured white adipocytes. Pathway activation is indicated by elevated phosphorylation of protein kinase A (PKA), PKA substrates (at the RRxpS/T consensus site), hormone sensitive lipase (HSL) perilipin (PLIN1), cyclic AMP response element binding protein (CREB) and extracellular signal regulated kinase (ERK). Beta-tubulin (TUBB) was used as a loading control. Elevated levels of phosphorylation within the lipolysis pathway are absent when *Opn3* null adipocytes are stimulated with blue light. (D) *Opn4$^{cre}$* activation of Ai14 identifies, as expected, neurons within the retinal ganglion cell layer. (B, C) *Opn4$^{cre}$* does not mark any adipocytes (or any cells at all) within interscapular brown adipose tissue (E, BAT) or inguinal white adipose tissue (F, inWAT). This indicates that *Opn4* is not normally expressed in these tissues. (G) Core body temperature in control (gray trace) and *Opn4* null (blue trace) mice during a 4°C exposure of 180 minutes. The absence of *Opn4* does not change the thermogenesis response. This is in contrast to *Opn3* loss-of-function mutant mice, in which the thermogenesis response is diminished. (H) Immunoblot assessing phospho-660-HSL levels in blue light stimulated white adipocytes from control and *Opn4* null mice. The absence of OPN4 does not prevent light-dependent phospho-HSL induction. Beta-tubulin (TUBB) was used as a loading control.
